# Plasmacytoid dendritic cells are functionally exhausted while non-haematopoietic sources of type I interferon dominate human autoimmunity

**DOI:** 10.1101/502047

**Authors:** Antonios Psarras, Adewonuola Alase, Agne Antanaviciute, Ian M. Carr, Md Yuzaiful Md Yusof, Miriam Wittmann, Paul Emery, George C. Tsokos, Edward M. Vital

## Abstract

Autoimmune connective tissue diseases arise in a stepwise fashion from asymptomatic preclinical autoimmunity. Type I interferons (IFNs) have a crucial role in the progression to established autoimmune diseases such as systemic lupus erythematosus (SLE). However, their cellular source and regulation in disease initiation are unclear. The current paradigm suggests that plasmacytoid dendritic cells (pDCs) are activated in SLE contributing to excessive IFN production. Here, we show that in preclinical autoimmunity, established SLE, and primary Sjögren’s Syndrome, pDCs are not effector cells, but rather have lost their capacity for TLR-mediated IFN-α and TNF production and fail to induce T cell activation, independently of disease activity and blood IFN signature. In addition, pDCs present a transcriptional signature of cellular stress and senescence accompanied by increased telomere erosion. Instead, we demonstrate a marked enrichment of IFN signature in non-lesional skin in preclinical autoimmunity. In these individuals and SLE patients, type I IFNs were abundantly produced by keratinocytes in the absence of infiltrating leucocytes. These findings revise our understanding of the role of IFN in the initiation of human autoimmunity, with non-haematopoietic tissues perpetuating IFN responses, which in turn predict clinical disease. These data indicate potential therapeutic targets outside the conventional immune system for treatment and prevention.

---

SLE and other autoimmune connective tissues diseases are a heterogeneous group of conditions. The pathogenesis of these diseases is incompletely understood. The most universal immune abnormality is the presence of autoantibodies targeting nuclear antigens. Dysregulation of the type I interferon (IFN) axis also plays a fundamental role (1, 2). Many lupus susceptibility genes are related to the IFN pathway (3–7). However, IFN activity is more variable; 60 – 80% of SLE patients exhibit increased expression of interferon-stimulated genes (ISGs) in peripheral blood (8–11).

Autoimmune connective tissue diseases are now recognized to arise in a stepwise fashion from asymptomatic preclinical autoimmunity. Autoantibodies precede symptoms by years and are far more common than clinical autoimmune disease (12–16). Hence, autoantibody-positive individuals constitute an “At-Risk” population of whom a minority will develop clinical autoimmunity. A key determinant of progression from At-Risk to established clinical autoimmune disease is the level of IFN activity (12). In established SLE, IFN activity is associated with cutaneous involvement and predicts future flares and severity (8, 17–19). In the present study we therefore asked how IFN production is controlled and regulated at both the preclinical and established autoimmune disease stages.

Most haematopoietic and non-haematopoietic cells are capable of producing type I IFNs (IFN-α, -β, -κ, -ω, -ε) as a first line of defence against viral infections. Nevertheless, much previous research on IFN production has focussed on plasmacytoid dendritic cells (pDCs). pDCs from otherwise healthy individuals produce particularly large amounts of type I IFNs upon recognition of viral antigens via endosomal toll-like receptors TLR7 and TLR9 (20, 21). Engagement of TLRs within endosomal compartments with these ligands leads to activation of IRF7 and NF-κB pathways and eventually production of IFN-α and other pro-inflammatory cytokines (TNF, IL-6) (22). Apart from this secretory function, pDCs exhibit antigen-presentation properties inducing both immunogenic and tolerogenic T cell responses (23–28). Conversely, the IFN-α-producing capacity of pDCs has been shown to be impaired in melanoma and ovarian cancer; tumour-infiltrating pDCs do not produce IFN-α but their presence actually promotes tumour growth (29–31). Additionally, hepatitis B virus can interfere with the TLR9 pathway by blocking MyD88-IRAK4 signalling and Sendai virus by targeting IRF7, while lymphocytic choriomeningitis virus (LCMV) compromises the capacity of pDCs to secrete type I IFNs. (32–34).

In the context of autoimmunity, endogenous nucleic acids forming immune complexes with autoantibodies have been proposed as a stimulus for pDC activation (35–38). While it is therefore natural to assume that pDCs are dominant producers of IFN-α in SLE, in fact the existing literature is complex and contradictory. Previous studies have reported both higher and lower numbers of pDCs in blood (39, 40). Unsorted PBMCs from SLE patients have been shown to produce lower levels of IFN-α in response to TLR9 stimulation, while other studies reported enhanced TLR7-mediated IFN-α production by pDCs of SLE patients (41, 42). It is not clear whether any alteration in pDC phenotype is the result of chronic inflammation or therapy, nor what underlying mechanism determines their function in early stages of human autoimmunity.

In order to resolve these contradictions, we assessed pDC phenotype, function and transcriptomic profile in At-Risk individuals who were therapy-naïve and did not have tissue inflammation, as well as established SLE and primary Sjögren’s Syndrome. We show that pDC numbers, TLR-mediated cytokine production and T cell-activating capacity are decreased prior to the onset of SLE and are unrelated to type I IFN activity. Instead, pDCs present a transcriptomic profile of cellular stress and senescence. In contrast, we show that non-haematopoietic tissue resident cells in the skin are not passive targets, but actively contribute to the aberrant type I IFN production seen in both preclinical and established autoimmunity. These findings provide unique insights into the source and regulation of type I IFNs in human autoimmune disease.

## RESULTS

### Circulating pDC numbers are decreased in preclinical autoimmunity and SLE

Peripheral blood pDCs were enumerated and immunophenotyped from freshly isolated peripheral blood mononuclear cells (PBMCs) using flow cytometry. We analyzed samples from At-Risk individuals (defined by ANA+, ≤1 clinical criterion for SLE, symptom duration <12 months and treatment-naïve; n = 64), patients with SLE (n = 81) and primary Sjögren’s Syndrome (pSS; n = 21) as well as age- and sex-matched healthy controls (n = 37). Clinical characteristics and treatment of SLE patients can be seen in **Supplemental Table 1**. pDCs were gated as CD3^−^CD19^−^CD14^−^CD56^−^CD11c^−^HLA-DR^+^CD123^+^CD303^+^ cells (**Figure 1A**). The average percentage of pDCs in PBMCs was significantly decreased in patients with SLE and pSS in comparison with healthy controls, a finding which was also observed in treatment-naïve At-Risk individuals (**Figure 1B**).

**Figure 1.**
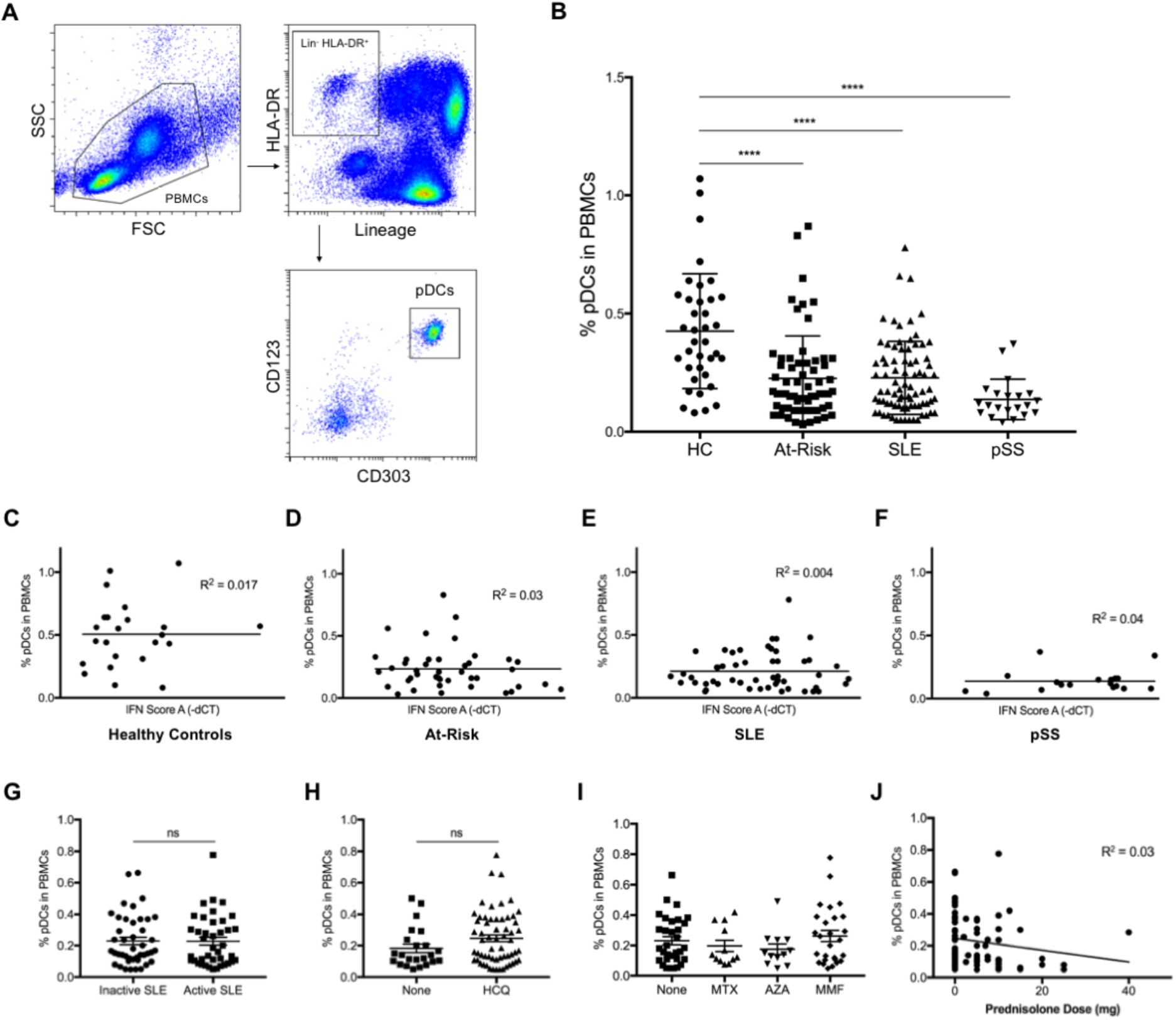
The percentage of pDCs in peripheral blood is decreased in At-Risk individuals and patients with SLE or pSS independently of IFN status, disease activity and treatment. **(A)** Gating strategy to identify pDC population within PBMCs: pDCs are characterized by the lack of expression of lineage markers (CD3, CD19, CD56, CD14, CD11c), intermediate to high expression of HLA-DR, high expression of CD123 (IL-3R) and CD303 (BDCA-2). **(B)** Average percentage of pDCs in PBMCs of age- and sex-matched healthy controls (HC; n = 37), At-Risk individuals (At-Risk; n = 64), patients with systemic lupus erythematosus (SLE; n = 81) and primary Sjögren’s Syndrome (pSS; n = 21). **(C-F)** Association between the percentage of pDCs in PBMCs and type I IFN activity in blood (IFN Score A) in HC, At-Risk, SLE, and pSS. **(G)** Percentage of pDCs in PbMcs in SLE patients with inactive and active disease. **(H)** Percentage of pDCs in PBMCs in SLE patients treated with or without hydroxychloroquine (HCQ). **(I)** Percentage of pDCs in PBMCs in SLE patients treated with other immunosuppressants (MTX, methotrexate; AZA, azathioprine; MMF, mycophenolate mofetil). **(J)** Association between the percentage of pDCs in PBMCs and the dose of prednisolone in patients with SLE. Data are represented as mean ± SEM. *ns* = not significant; \*\*\*\**p* < 0.0001. 2-way ANOVA (**B**), nonlinear regression (**C-F** and **J**), unpaired 2-tailed *t* test (**G-I**).

To determine whether the reduction of circulating pDCs was associated with blood ISG expression and other clinical features, we evaluated the expression of a previously validated IFN score (8) (IFN Score A) in PBMCs using TaqMan in all sample groups described above. Increased type I IFN activity as assessed by IFN Score A was observed in patients with SLE and pSS as well as At-Risk individuals compared to healthy controls, but the reduction of peripheral blood pDCs described above was not associated with the higher expression of IFN Score A (**Figure 1C-F**). Although IFN Score A was associated with increased number of extractable nuclear antigen (ENA) antibodies, no association was found between the percentage of pDCs and either IFN Score A or ENA antibodies in any of the sample groups (**Supplemental Figure 1A-B**). Additionally, in SLE patients the reduction of circulating pDCs was independent of disease activity (**Figure 1G**), treatment with hydroxychloroquine (**Figure 1H**), other immunosuppressants (**Figure 1I**) or prednisolone (**Figure 1J**). Apart from that, the reduction of circulating pDCs in At-Risk individuals, patients with SLE and pSS was not associated with the total lymphocyte count, which is commonly low in SLE patients (**Supplemental Figure 1C-E**).

pDCs from healthy controls, At-Risk individuals and SLE patients were analyzed for the surface expression of multiple molecules known to be important in regulating their immune functions (**Supplemental Figure 2A-F**). pDCs in SLE patients showed no statistically significant difference in the expression of HLA-DR or BDCA-2 (CD303). On the other hand, CD123 (IL-3R) and ILT2 (CD85j) were found to be upregulated on pDCs of SLE patients compared to healthy controls (*P* < 0.001). Interestingly, CD317 (BST2; tetherin), a molecule known to be induced by type I IFNs, also presented higher expression on pDCs of SLE patients (*P* < 0.05); however, its ligand ILT7 (CD85g) appeared to be downregulated on pDCs of SLE patients (*P* < 0.05).

**Figure 2.**
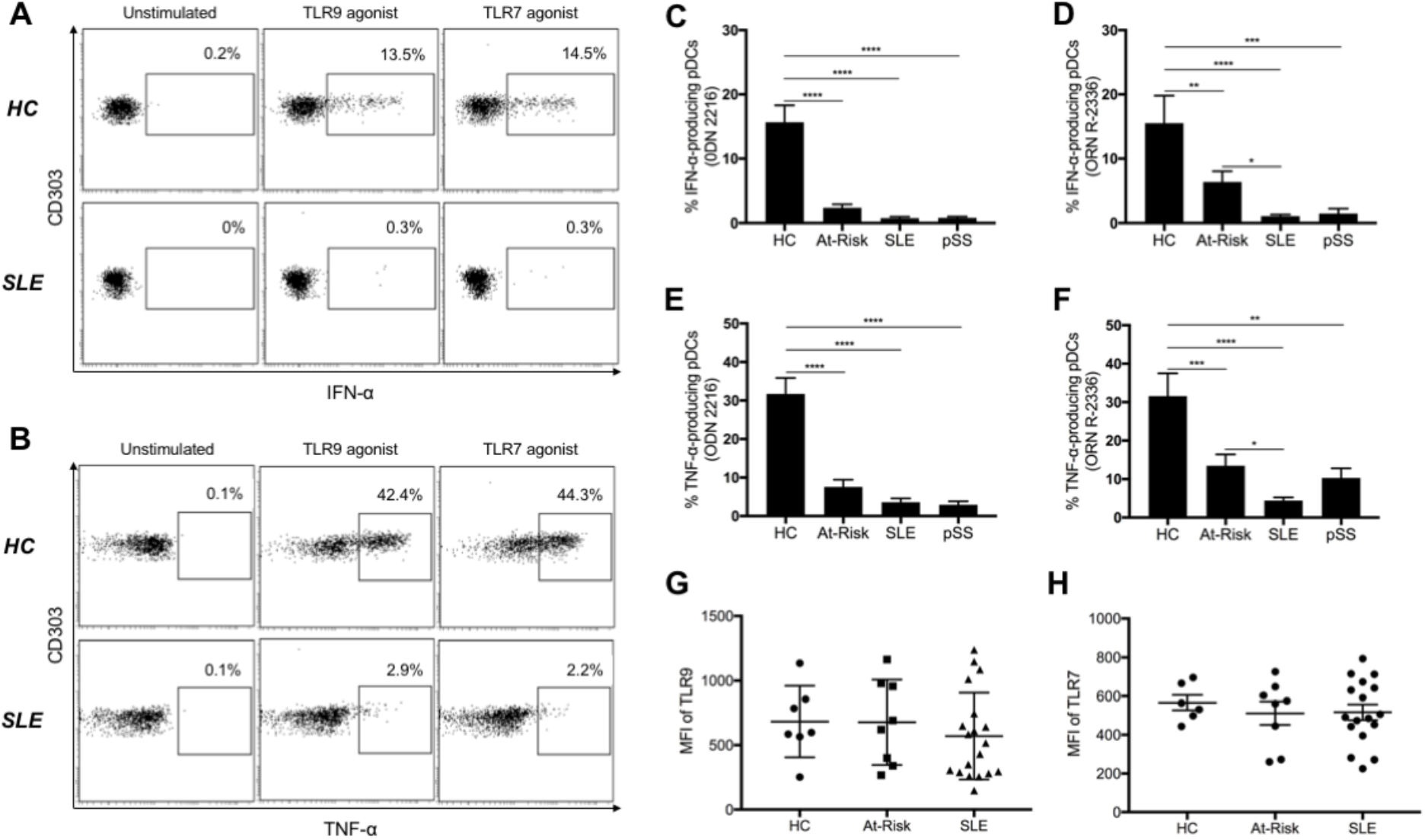
pDCs from At-Risk, SLE and pSS patients produce less IFN-α and TNF after stimulation with synthetic TLR agonists. **(A-B)** Freshly isolated PBMCs were cultured in the absence or presence of TLR9 (ODN 2216) or TLR7 (ORN R-2336) agonists for 6 hours, then IFN-α and TNF production by pDCs was measured using intracellular staining. Results shown are representative of a healthy control (HC) and a patient with SLE. Average percentage of IFN-α produced by TLR9-stimulated **(C)** and TLR7-stimulated **(D)** pDCs in HC (n = 14), At-Risk (n = 26), SLE (n = 40) and pSS (n = 7) patients. Average percentage of TNF produced by TLR9-stimulated **(E)** and TLR7-stimulated **(F)** pDCs in HC (n = 14), At-Risk (n = 26), SLE (n = 40) and pSS (n = 7) patients. **(G-H)** Intracellular expression of TLR9 and TLR7 was measured using flow cytometry in HC (n = 7), At-Risk (n = 8) and SLE (n = 19) patients. Data are represented as mean ± SEM. \**P* <0.05; \*\**P* < 0.01; \*\*\**P* < 0.001; \*\*\*\**P* < 0.0001. 2-way ANOVA (**C-H**).

### TLR-stimulated pDCs produce less cytokines in preclinical autoimmunity and SLE

The production of IFN-α and other pro-inflammatory cytokines (e.g. TNF) in response to TLR-mediated stimulation is the hallmark of pDC function. To evaluate the capacity of cytokine production by pDCs, we stimulated freshly isolated PBMCs from At-Risk individuals (n = 26), patients with established SLE (n = 40) and pSS (n = 7) alongside healthy controls (n = 14) for 6 hours with TLR9 (ODN 2216) or TLR7 (ORN R-2336) agonists. We measured both IFN-α and TNF produced by CD3^−^CD19^−^CD14^−^CD56^−^CD11c^−^HLA-DR^+^CD123^+^CD303^+^ pDCs using intracellular staining. No IFN-α and/or TNF production by pDCs was detected in any of the samples without external stimulation. While pDCs from healthy controls produced large amounts of IFN-α in response to TLR9 or TLR7 agonists, pDCs from SLE patients showed little or no cytokine production (**Figure 2A**). Furthermore, TLR9- and TLR7-mediated IFN-α production was diminished in pDCs from patients with pSS similarly to SLE (**Figure 2C-D**). Although TLR9-mediated IFN-α production was similarly reduced in At-Risk individuals, their pDCs seemed to partially maintain some TLR7-mediated IFN-α production (**Figure 2D**). TLR9- and TLR7-mediated TNF production was also significantly decreased in pDCs from patients with SLE and pSS compared to healthy controls (**Figure 2B**), whilst pDCs from At-Risk individuals showed the same trend as for IFN-α production, partially maintaining some TLR7-mediated TNF production (**Figure 2E-F**).

We next evaluated whether there was any association between IFN-α production by pDCs and overall blood type I IFN activity as measured by IFN Score A. We found no association between the levels of TLR-mediated IFN-α production and the level of IFN Score A in patients with SLE and pSS as well as At-Risk individuals (**Supplemental Figure 3A-D**). To confirm that these findings were not due to differences in TLR expression, we measured the expression levels of both TLR9 and TLR7 using flow cytometry. pDCs from At-Risk individuals and SLE patients showed similar expression levels of both receptors compared to those of healthy controls (**Figure 2G-H**).

**Figure 3.**
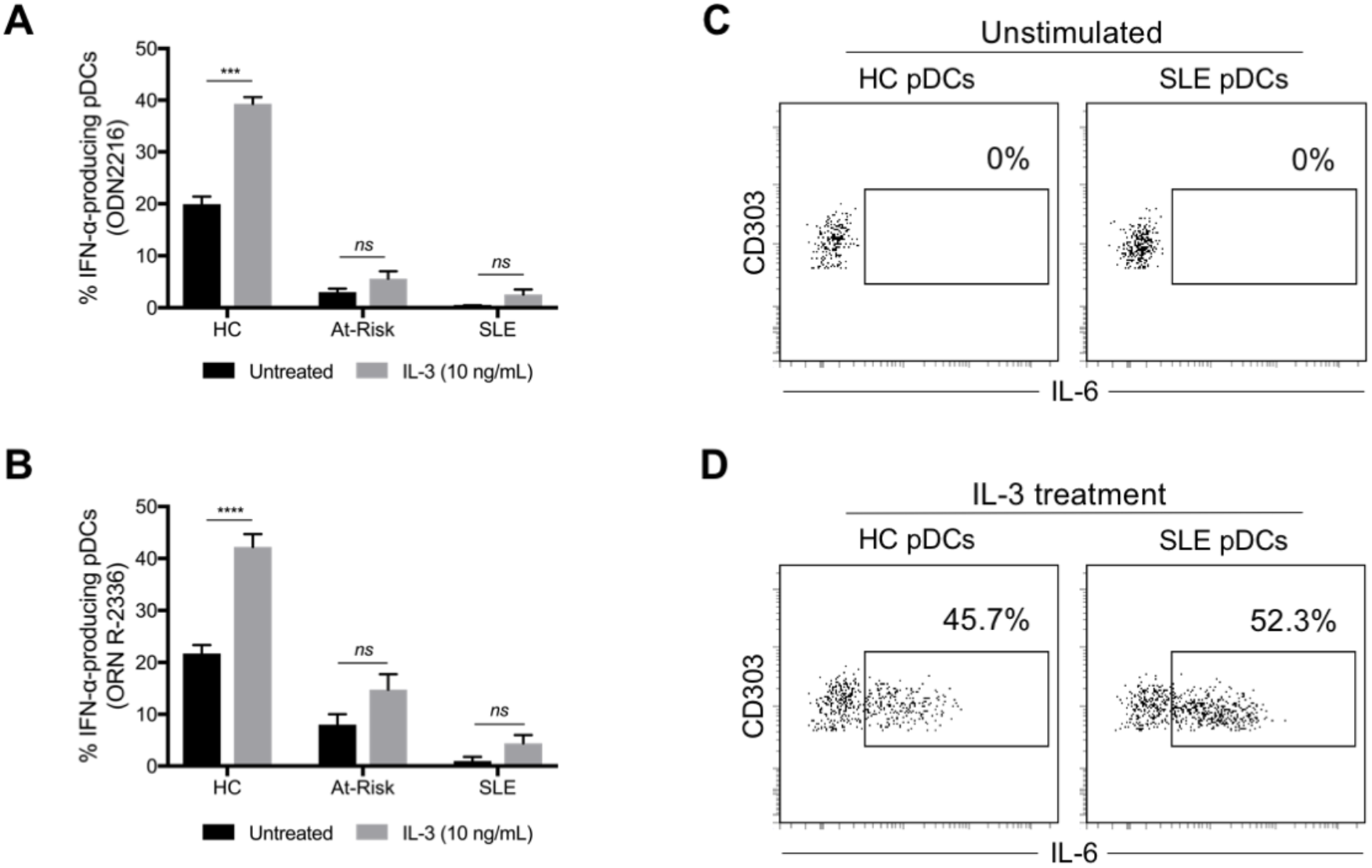
IL-3 triggers TLR-independent production of IL-6 by pDCs. PBMCs from healthy controls (HC; n = 6), At-Risk individuals (At-Risk; n = 4) and SLE patients (n = 7) were cultured for 18 hours in the absence or presence of IL-3 (10 ng/mL). The cells were then stimulated by TLR9 (ODN 2216) or TLR7 (ORN R-2336) agonists for 6 additional hours. The production of cytokines was measured by intracellular staining. **(A)** IL-3 significantly enhanced TLR9-mediated IFN-α production by pDCs of healthy controls (*P* < 0.001); this effect was not seen in pDCs of At-Risk (*P* = 0.3) and SLE (*P* = 0.4) patients. **(B)** IL-3 significantly enhanced TLR7-mediated IFN-α production by pDCs of healthy controls (*P* < 0.0001); this effect was not that prominent in pDCs of At-Risk (*P* = 0.09) and SLE (*P* = 0.6) patients. **(C and D)** Treatment with IL-3 (10 ng/mL) induced the production of IL-6 by pDCs of both healthy controls and SLE patients without exogenous TLR stimulation. The production of IL-6 was detected by intracellular staining. Data are represented as mean ± SEM. \*\*\**P* < 0.001; \*\*\**P* < 0.0001; *ns* = not significant. 2-way ANOVA (**A**).

Interestingly, while culturing PBMCs, we observed that, a population within monocytes was characterized by no expression of HLA-DR but positive expression of CD303 (BDCA-2), which was previously thought to be a pDC-specific marker and used in immunohistochemistry. These cells showed no response to TLR stimulation, as neither IFN-α nor TNF production was detected (**Supplemental Figure 3E-F**).

### IL-3 triggers TLR-independent production of IL-6 by pDCs

IL-3 is known to maintain pDC survival *in vitro* and to enhance IFN-α production upon TLR-mediated stimulation (43, 44). We confirmed that pre-treatment for 24 hours with IL-3 amplified IFN-α production by both TLR9- and TLR7-stimulated pDCs from healthy controls (n = 6). However, a statistically significant increase in IFN-α production was not seen in pDCs of At-Risk individuals (n = 4) and SLE patients (n = 7) (**Figure 3A-B**). Furthermore, we discovered that IL-3 triggered the spontaneous production of IL-6 by pDCs without any exogenous TLR-mediated stimulation. In contrast to the defective IFN-α and TNF production in pDCs from SLE patients we described above, this TLR-independent IL-6 production was not impaired in pDCs of SLE patients (**Figure 3C-D**).

### pDCs from SLE patients display decreased ability to induce T cell activation

Although pDCs possess antigen-presentation properties and can trigger T cell responses, little is known about the capacity of pDCs in SLE to induce T cell proliferation and activation. We co-cultured freshly isolated pDCs from patients with active SLE and healthy controls with CellTrace Violet-labelled allogeneic naïve CD4^+^ T cells in the presence of low ratio anti-CD3/CD28 beads for 5 days (**Figure 4A**).

**Figure 4.**
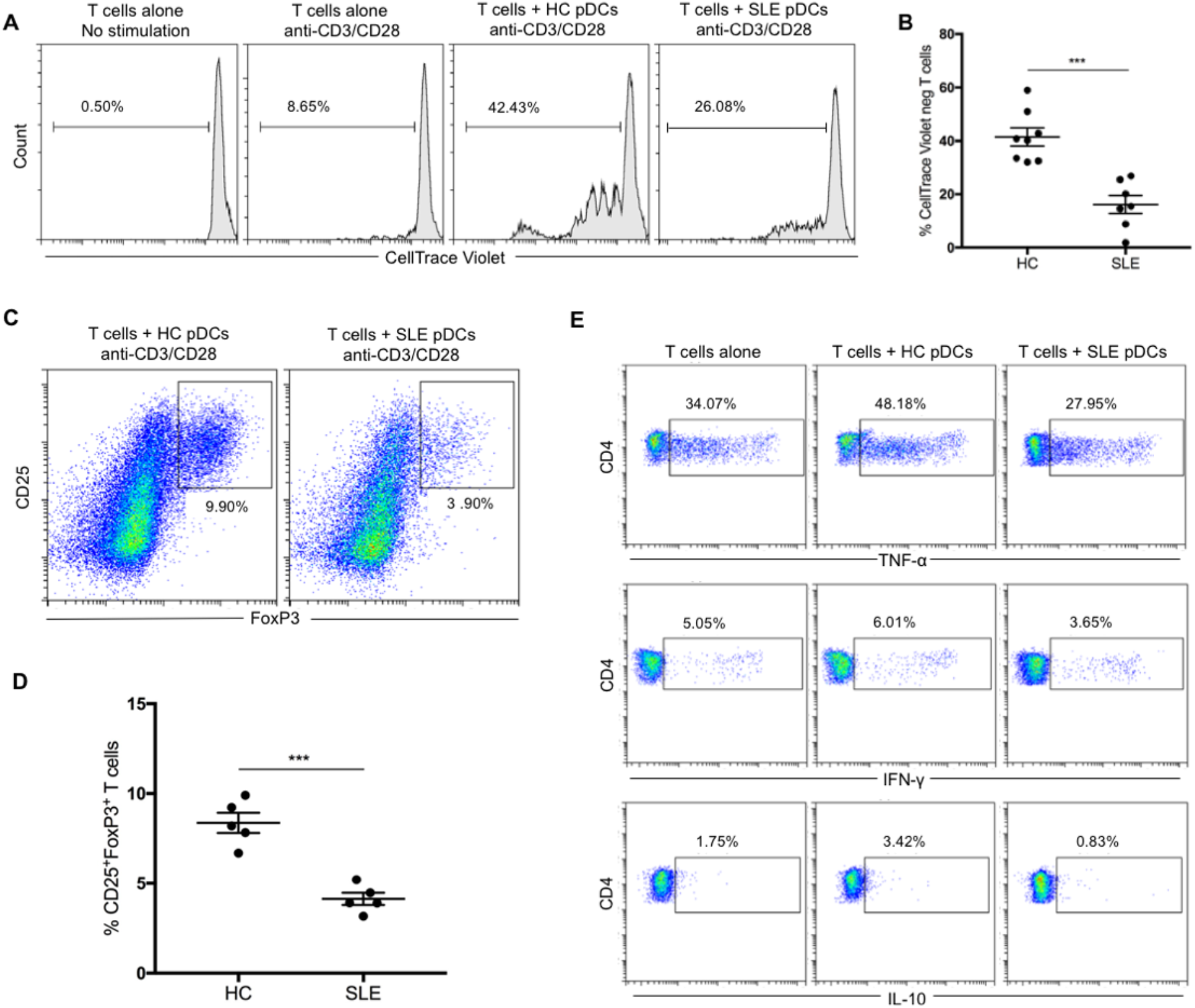
pDCs from SLE patients display decreased ability to induce T cell proliferation and activation. **(A)** Allogeneic naïve CD4^+^ T cells were labeled with CellTrace Violet and cultured alone or with pDCs purified from healthy controls (HC) or patients with active SLE for 5 days in the presence of anti-CD3/CD38 beads at ratio 2:1 to avoid excessive T cell activation and expansion. T cell proliferation was analyzed by flow cytometry based on CellTrace Violet dilution. One representative experiment is shown out of four independent experiments. **(B)** Average percentage of proliferated CD4^+^ T cells co-cultured with pDCs from healthy controls (n = 8) and SLE patients (n=7). **(C)** Induction of CD4^+^CD25^high^FoxP3^+^ T cells from naïve CD4^+^ T cells co-cultured for 5 days with pDCs from healthy controls or SLE patients in the presence of anti-CD3/CD28 beads at ratio 2:1. One representative experiment is shown out of three independent experiments. **(D)** Percentage of CD4^+^CD25^high^FoxP3^+^ T cells derived from the co-culture with pDCs from healthy controls (n = 5) and SLE patients (n = 5). **(E)** Allogeneic naïve CD4^+^ T cells were cultured alone or with pDCs from healthy controls or SLE patients for 5 days in the presence of anti-CD3/CD38 beads at ratio 2:1. On the fifth day, the cells were stimulated with PMA/Ionomycin and the production of TNF, IFN-γ and IL-10 by CD4^+^ T cells was measured by intracellular staining. One representative experiment is shown out of three independent experiments. Data are represented as mean ± SEM. \*\*\**P* < 0.001. Unpaired 2-tailed *t* test (**B** and **D**).

Although pDCs from both groups induced T cell proliferation, pDCs from SLE patients were substantially less efficient in inducing T cell proliferation (**Figure 4B**). pDCs are also known to trigger the induction of FoxP3^+^ T cells. Following the same protocol as above, we found that fewer CD25^high^FoxP3^+^ cells were generated from naïve CD4^+^ T cells after 5 days of co-culturing with pDCs from SLE patients in comparison with pDCs from healthy controls (**Figure 4C-D**).

To investigate the ability of pDCs to trigger cytokine production by T cells, we first co-cultured pDCs from patients with active SLE and healthy controls with allogeneic naïve CD4^+^ T cells in the presence of low ratio anti-CD3/CD28 beads for 5 days before PMA/Ionomycin were added in the last 5 hours of the culture. In comparison with T cells alone, pDCs from healthy controls enhanced the production of TNF (34.07% vs. 48.18%), IFN-γ (5.05% vs. 6.01%) and IL-10 (1.75% vs. 3.42%) from the co-cultured T cells (**Figure 4E**). However, pDCs from SLE patients suppressed the production of all cytokines measured; TNF (34.07% vs. 27.95%), IFN-γ (5.05% vs. 3.65%) and IL-10 (1.75% vs. 0.83%). In summary, pDCs from SLE patients exhibit decreased capacity for triggering T cell proliferation and activation, whilst they actively inhibited cytokine production by T cells.

### pDCs from IFN^low^ and IFN^high^ SLE patients present distinct transcriptomic profiles

To investigate disease-associated transcriptional changes in pDCs, we purified pDCs from healthy controls (n = 8), At-Risk individuals (n = 4) and SLE patients (n = 13) by negative selection and we then sorted the cells to achieve purity >99% based on CD304 (BDCA-4) expression. We sequenced the RNA extracted from sorted pDCs using Smart-seq2 for sensitive full-length transcriptomic profiling.

A major source of variability amongst the samples was due to the expression of ISGs. In order to control for this variability, we therefore first scored each sample based on the expression profile of a core set of ISGs (IFN Score). The expression level of IFN Score in pDCs from healthy controls (95% CI) was used to assign each sample to IFN^low^ or IFN^high^ subgroups (**Figure 5A**). As expected, pDCs from SLE patients were characterized by a range of IFN Scores, but overall exhibited a higher IFN Score than pDCs from healthy controls and At-Risk individuals (**Figure 5B**). pDCs from most At-Risk individuals (3/4) presented a higher IFN Score compared to pDCs from healthy controls and they were assigned to the IFN^high^ subgroup. Common ISGs (*MX1*, *XAF1*, *IFI44*, *RSAD2*) were found to be upregulated in the majority of pDCs in IFN^high^ SLE patients and At-Risk individuals, whilst pDCs of IFN^low^ SLE patients showed similar expression levels to those of healthy controls (**Figure 5C**).

**Figure 5.**
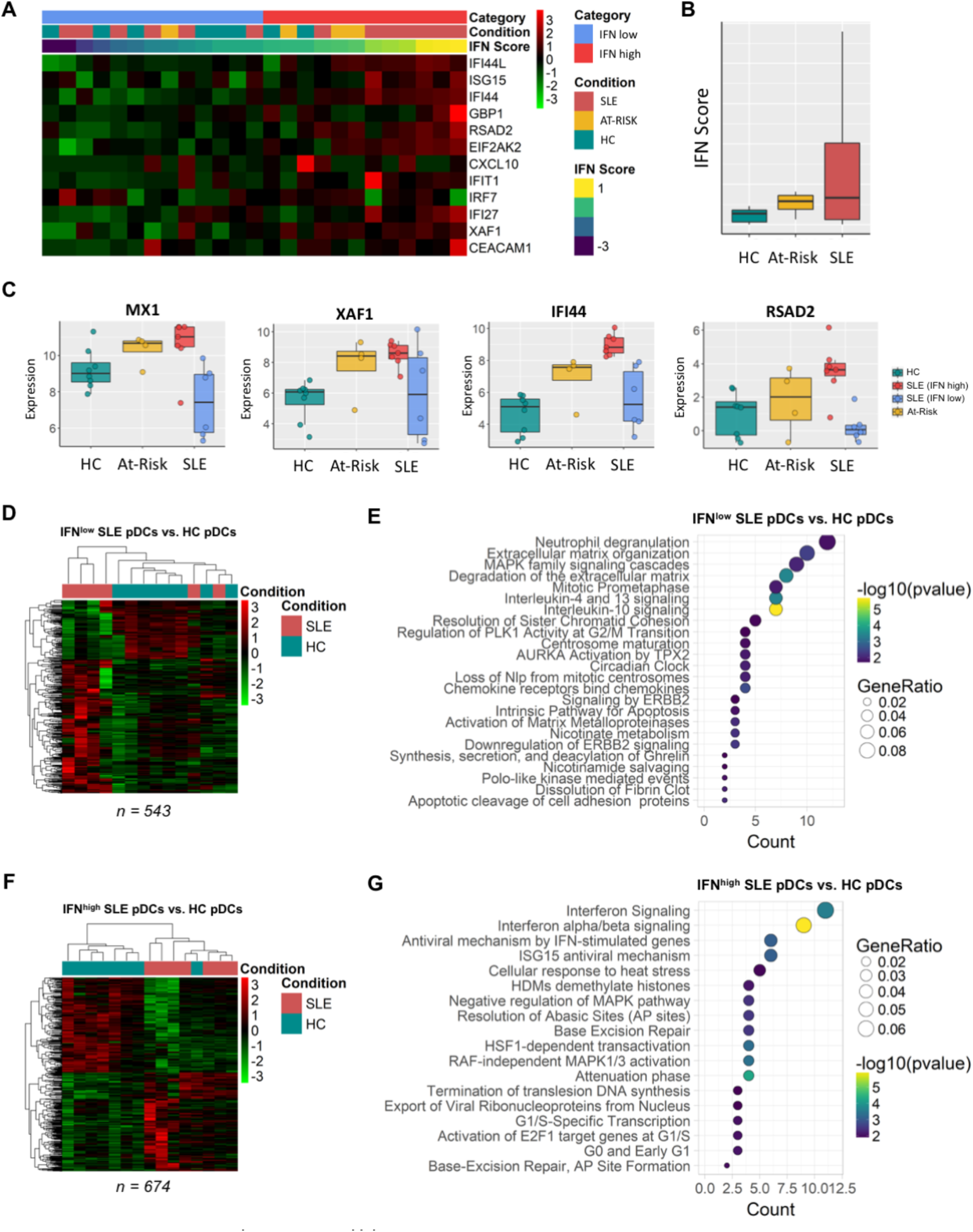
pDCs from IFN^low^ and IFN^high^ SLE patients display distinct transcriptomic profiles. **(A)** Sorted pDCs from HC (n = 7), At-Risk (n = 4) and SLE (n = 13) were classified according to the expression level of the IFN Score described. **(B)** Average expression level of IFN Score measured in samples described in **(A)**. **(C)** Expression level of representative ISGs in sorted pDCs from sample groups described in **(A)**. **(D)** Differentially expressed transcripts (n = 543) in IFN^low^ SLE pDCs compared to HC pDCs. **(E)** Reactome Pathway Enrichment in differentially expressed genes of IFN^low^ SLE pDCs. **(F)** Differentially expressed transcripts (n = 674) in IFN^high^ SLE pDCs compared to HC pDCs. **(G)** Reactome Pathway Enrichment in differentially expressed genes of IFN^high^ SLE pDCs.

Following this classification, we first sought to investigate changes in gene expression profiles of pDCs associated with each IFN^low^ or IFN^high^ subgroup compared to those of healthy controls, then subsequently to identify the differentially expressed in genes in SLE patients common to both IFN subgroups.

The analysis of IFN^low^ patients revealed 543 transcripts that were significantly (FDR < 5%) differentially expressed (**Figure 5D**), which were particularly enriched for IL-4 and IL-13 signalling, IL-10 signalling, cell migration and pathogen interaction pathways, amongst others (**Figure 5E and Supplemental Figure 4A**). Amongst the upregulated genes were chemokines, for instance *CXCL3*, *CXCL2* and *CXCL16* (**Supplemental Figure 5**). A detailed table of the top differentially expressed genes in pDCs of IFN^low^ SLE patients can be found in **Supplemental Table 2**.

In IFN^high^ SLE patients, we found 674 transcripts that were significantly (FDR < 5%) differentially expressed (**Figure 5F**). Unsurprisingly, these genes were found to be heavily enriched for IFN-response related pathways (**Figure 5G and Supplemental Figure 4B**), but also pathways related to DNA repair and MAPK signalling. Several phosphatases known to dephosphorylate MAP kinases (*DUSP1*, *DUSP2*, *DUSP5* and *DUSP8*), transcriptional repressors associated with cell differentiation (*HESX1*, *ETV3*) and NF-κB inhibitors (*NFKBIA*, *NFKBID*) were found to be upregulated in IFN^high^ SLE patients (**Supplemental Figure 5**). A detailed table of the top differentially expressed genes in pDCs of IFN^high^ SLE patients can be found in **Supplemental Table 3**.

We also evaluated the expression levels of multiple TLRs in the pDCs of all samples (**Supplemental Figure 6**). As expected, pDCs from healthy controls, At-Risk individuals and SLE patients strongly expressed TLR9 and TLR7; however, no differences were found among the different groups regardless of IFN activity, as also confirmed by the intracellular expression of TLR9 and TLR7 at protein level (**Figure 2G-H**). Lower expression of TLR1, TLR6, and TLR10 was observed in pDCs with no difference among the groups, but no expression of TLR2, TLR3, TLR4, TLR5, and TLR8 was found in any of the samples. Additionally, no transcripts for the 14 distinct subtypes of IFN-αlpha were detected in the pDCs of any sample. No other type I IFN transcripts (IFN-beta, IFN-kappa, IFN-omega) as well as transcripts for type III IFNs (IFN-lambda) were found to be expressed (**Supplemental Figure 7**). No significant differences were detected in expression of chemokines and their receptors apart from the ones described in pDCs of IFN^low^ or IFN^high^ SLE patients (**Supplemental Figure 8 and 9**).

**Figure 6.**
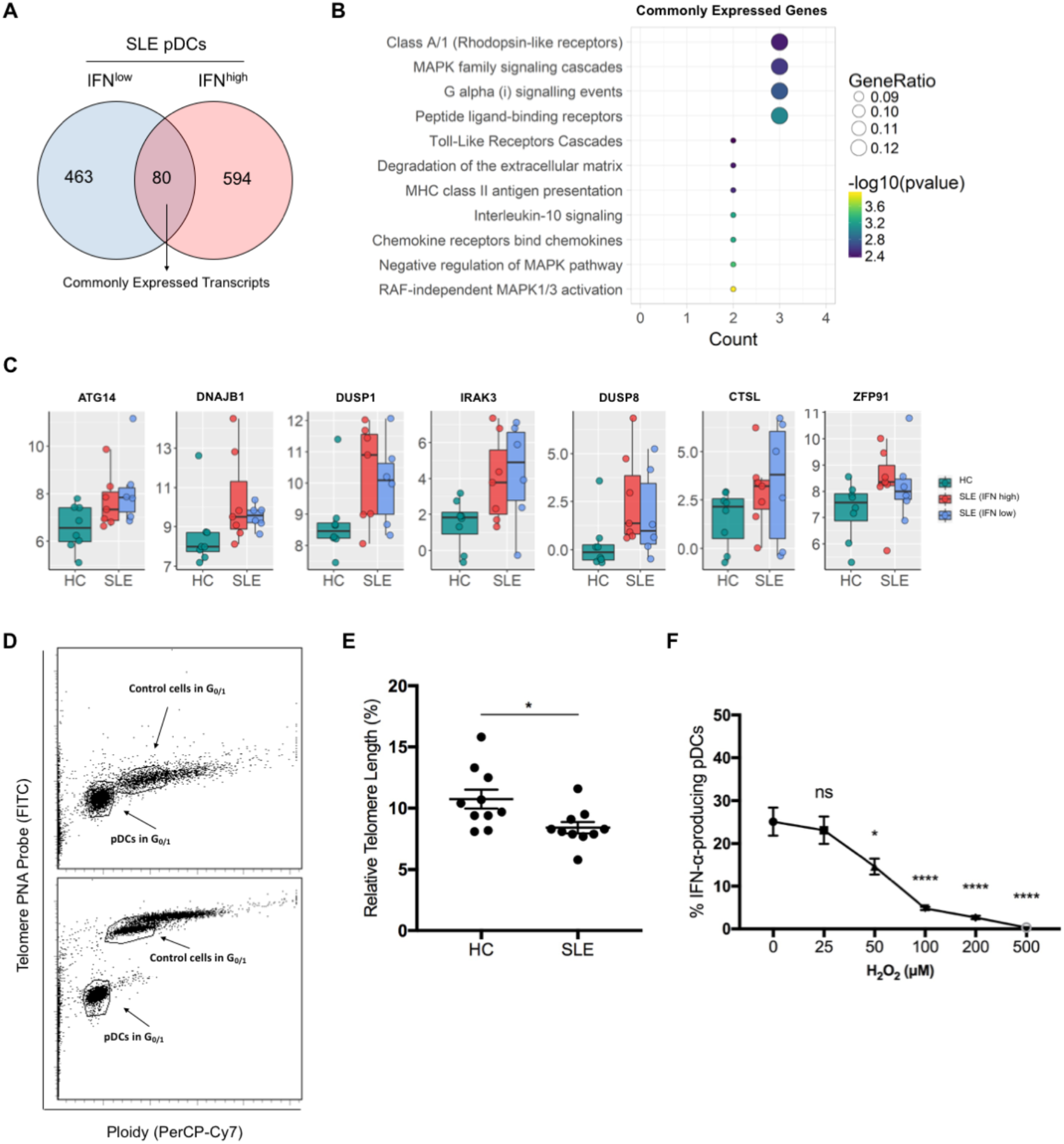
pDCs from SLE patients present transcriptional and phenotypic features of immune senescence. **(A)** Venn diagram showing the number of differentially expressed transcripts (n = 80) common to both IFN^low^ and IFN^high^ pDCs from SLE patients compared to pDCs from HC. **(B)** Reactome Pathway Enrichment in DEGs in differentially expressed genes in IFN^low^ and IFN^high^ pDCs from SLE patients shown in **(A)**. **(C)** Expression level of representative genes differentially expressed in both IFN^low^ and IFN^high^ pDCs from SLE patients in comparison with pDCs from HC. (Purified pDCs from freshly isolated PBMCs were hybridized without **(D; upper)** or with **(D; lower)** telomere PNA probe. Gates were set in G_0/1_ phase for both sample cells (pDCs) and tetraploid control cells (1301 cell line). **(E)** Determination of the relative telomere length as the ratio between the telomere signal of pDCs purified from HC (n = 10) and SLE (n = 10) patients and the control cells (1301 cell line) with correction for the DNA index of G_0/1_ cells. **(F)** Freshly isolated PBMCs from healthy donors (n = 4) were exposed to H_2_O_2_ (0 – 500 μM) for 15 minutes. After H_2_O_2_ exposure, cells were washed thoroughly and resuspended in culture medium before they were stimulated with 2μM ODN 2216 for 6 hours. The production of IFN-α by pDCs was measured in viable cells by intracellular staining. Data are represented as mean ± SEM. *ns* = not significant; \**P* < 0.05; \*\*\*\**P* < 0.001. Unpaired 2-tailed *t* test (**E**), 1-way ANOVA (**F**).

### pDCs from SLE patients present transcriptional and phenotypic features of immune senescence

*In vitro* functional assays demonstrated that the decreased secretory function upon TLR stimulation was universally observed in pDCs of SLE patients, independently of the IFN activity in their PBMCs (**Figure 2G**). To investigate which biological pathways contribute to this defective phenotype, we therefore studied the transcripts differentially expressed in both pDCs of IFN^low^ and IFN^high^ SLE patients compared to those of healthy controls. Little overlap between differentially expressed genes in pDCs of IFN^low^ and IFN^high^ SLE patients was detected (**Figure 6A**). Reactome Pathway Enrichment also showed that biological processes related to MAPK family signalling, TLR signalling, IL-10 signalling and chemotaxis were significantly enriched (**Figure 6B and Supplemental Figure 4C**). Amongst the 80 shared transcripts, there were upregulated genes involved in cellular senescence and stress (*ATG14*, *ATP7A*, *DNAJB1*), protein degradation in lysosomes (*CTSL*), negative regulation of TLR signalling (*IRAK3*), negative regulation of MAPK signalling (*DUSP1*, *DUSP8*) and negative regulation of non-canonical NF-κB pathway (*ZFP91*), which are all known to inhibit the production of type I IFNs and other pro-inflammatory cytokines (**Figure 6B-C**). Moreover, the shared transcripts included upregulated genes for *CXCL2* and *CCL19* (**Supplemental Figure 5**). A detailed table of the genes commonly differentially expressed in pDCs of both IFN^low^ and IFN^high^ SLE patients can be found in **Supplemental Table 4**.

Increased telomere erosion is known to be related to cellular senescence, a feature has been found in other immune cells of patients with SLE but has not previously been described in pDCs (45). To address this question, we purified pDCs from healthy controls alongside SLE patients, which were then hybridized with telomere PNA probe before they were analyzed by flow cytometry. The relative telomere length was calculated as the ratio between the telomere signal of pDCs and the tetraploid control cells (1301 cell line) with correction for the DNA index of G_0/1_ cells (**Figure 6D**). The analysis confirmed that pDCs from SLE patients had shorter telomere length compared to pDCs from age- and sex-matched healthy controls (**Figure 6E**).

RNA-sequencing data suggested that genes related to cellular stress are amongst the 80 commonly shared transcripts between pDCs of IFN^low^ and IFN^high^ SLE patients compared to pDCs of healthy controls. Thus, we next sought to investigate the effect of oxidative stress on type I IFN production in TLR-stimulated pDCs. Freshly isolated PBMCs from healthy donors were exposed to increasing concentrations of H_2_O_2_ (0 − 500 μM) for 15 minutes before they were stimulated with ODN 2216 and IFN-α production was measured in viable cells by intracellular staining. We observed that oxidative stress –even at low concentrations of H_2_O_2_- negatively regulated TLR-mediated responses in pDCs leading to a gradual loss of their ability to produce IFN-α (**Figure 6F**).

### IFN activity is increased in non-lesional skin in preclinical autoimmunity

Since professional IFN-α-producing cells such as pDCs appeared immunosenescent in SLE, the source of the aberrant type I IFN production seen in patients had yet to be identified. We compared the level of expression of IFN Score A in blood with disease activity in the two most common organ manifestations, defining active disease as BILAG-2004 A or B and inactive disease as BILAG-2004 C-E. We found that IFN Score A was associated with mucocutaneous disease activity (Fold Difference 2.24 (95% CI 1.16 - 4.34); *P* = 0.017), but not with musculoskeletal disease (Fold Difference 0.97 (95% CI 0.44 - 2.09); *P* = 0.927) (**Figure 7A-B**).

**Figure 7.**
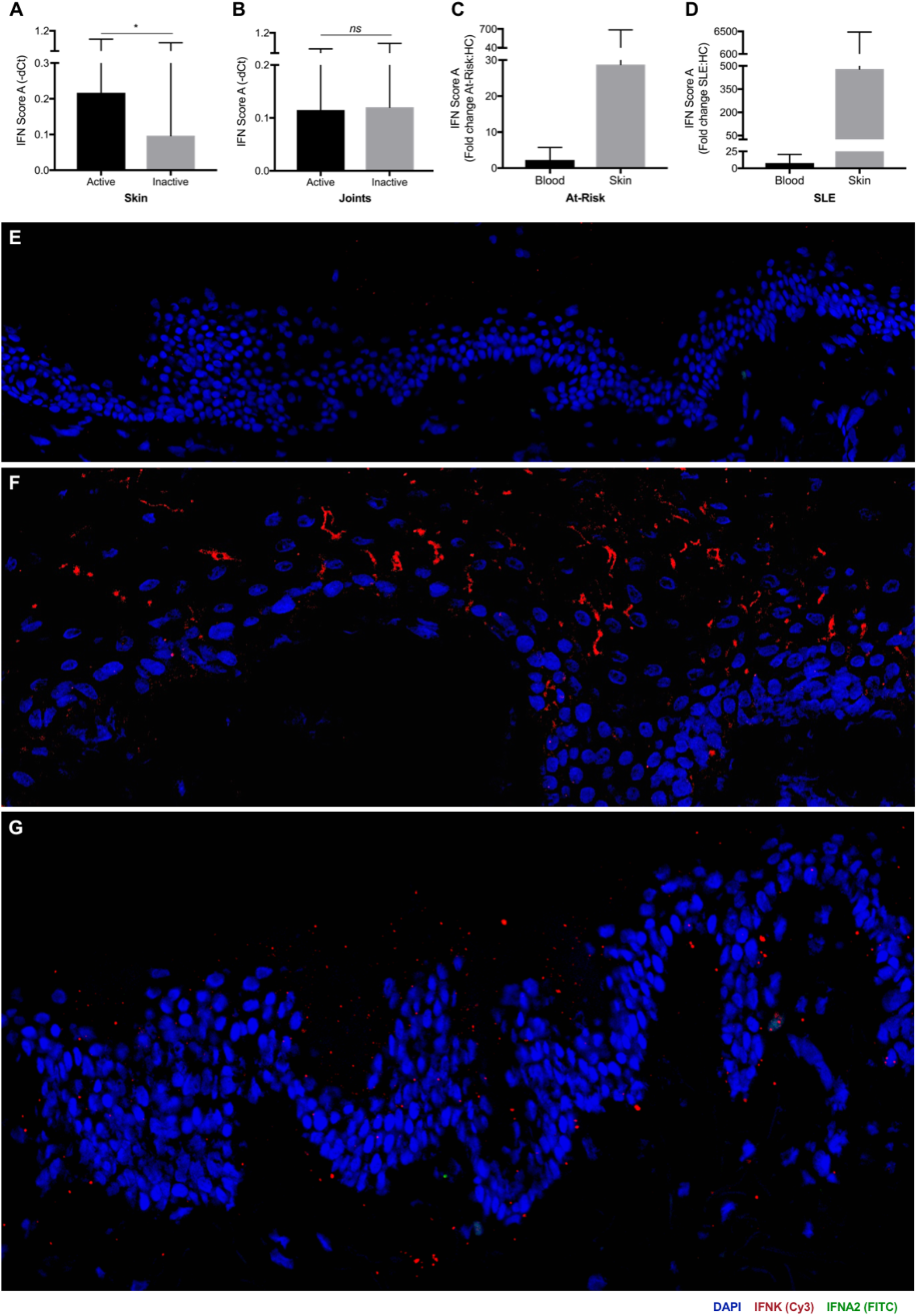
SLE patients and At-Risk individuals present diffuse expression of type I IFNs in epidermis. Association of IFN Score A with active and inactive mucocutaneous disease in SLE patients (**A**). Association of IFN Score A with active and inactive musculoskeletal disease in SLE patients (**B**). Fold increase in IFN Score A of At-Risk individuals (**C**) in blood (2.21; 1.37, 3.53) and skin (28.74; 1.29, 639.48) compared to healthy controls. Fold increase in IFN Score A of SLE patients (**D**) in blood (7.80; 4.75,12.80) and skin (479.33; 39.32, 5842.78) compared to healthy controls. Skin biopsies were hybridized using RNAscope *in situ* hybridization technology with custom-designed target probes for *IFNA2* and *IFNK*. Hybridization signals were amplified and detected using TSA Plus fluorescein (FITC) for *IFNA2* and TSA Plus Cyanine 3 (Cy3) for *IFNK*. Nuclei were highlighted using DAPI. Representative *in situ* hybridization images of: **(E)** healthy control, **(F)** IFN^high^ SLE patient with active skin lesion, **(G)** IFN^high^ At-Risk individual with no clinical or histopathological signs of inflammation. Data are represented as mean ± SEM. *ns* = not significant; \**P* < 0.05. Unpaired 2-tailed *t* test (**A** and **B**).

Next, we compared the fold increase in IFN Score A in blood and skin biopsies from At-Risk individuals and SLE patients compared to healthy controls (**Figure 7C-D**). We analyzed blood samples from 114 SLE patients, 105 At-Risk individuals, and 49 healthy controls; we also analyzed lesional skin biopsies from 10 SLE patients and non-lesional skin biopsies from 10 At-Risk individuals as well as skin biopsies from 6 healthy controls. In At-Risk individuals compared to healthy controls, mean (95% CI) fold increase in IFN Score A in blood was 2.21 (1.37, 3.53), while in non-lesional skin the fold increase was markedly higher at 28.74 (1.29, 639.48). The differential increase in ISG expression in blood and skin was even more extreme in SLE patients compared to healthy controls; in some SLE patients ISG expression in skin was more than 5,000 times higher than healthy controls. Mean (95% CI) fold increase was 7.80 (4.75,12.80) in blood compared to 479.33 (39.32, 5842.78) in skin.

### Patients with high IFN activity in blood present diffuse expression of type I IFNs in epidermis

Given this extreme elevation of ISG expression in skin compared to blood, we further analyzed skin biopsies from healthy controls (n = 4), SLE patients (n = 6) and At-Risk individuals (n = 4). Skin biopsies were obtained from active lesions of SLE patients, whilst skin biopsies from At-Risk individuals had no clinical or histopathological signs of inflammation. We performed *in situ* hybridization using RNAscope technology to visualise the direct expression of type I IFNs transcripts (*IFNK, IFNA2*) at a cellular level in all skin biopsies obtained.

As expected, skin biopsies from healthy controls with minimal IFN Score A in blood showed no expression of either *IFNK* or *IFNA2* (**Figure 7E**). In contrast, active skin lesions from SLE patients with high IFN Score A in blood demonstrated diffuse expression of *IFNK* in the epidermis (**Figure 7F**). Intriguingly, the epidermis of At-Risk individuals with high IFN Score A in blood was also characterized by diffuse expression of *IFNK*, although unlike SLE patients, there were no clinical or histopathological features of inflammation (**Figure 7G**). Regarding *IFNA2* expression, we detected expression in the dermis, possibly by fibroblasts as *IFNA2* signal was located within dense connective tissue. However, notably we did not see expression of *IFNK* or *IFNA2* in areas of leucocyte infiltration (**Supplemental Figure 10**). Collectively, these results indicate that the high IFN activity observed in both non-lesional and lesional skin, including at a preclinical phase, was not mediated by infiltrating haematopoietic immune cells. Instead, the *in situ* hybridisation suggested that non-haematopoietic cells, such as keratinocytes throughout histologically normal skin, were responsible for type I IFN production in scenarios where pDCs produced none.

### Keratinocytes from At-Risk and SLE patients present increased expression of IFNs in response to UV light and nucleic acids

To test the response of keratinocytes to a known environmental trigger for cutaneous inflammation, we measured the expression of type I IFN transcripts in non-lesional skin of a patient with clinically inactive SLE before and after UV provocation *in vivo*. In a biopsy obtained before UV provocation, there was low *IFNK* expression in the epidermis (**Figure 8A**). We then obtained a second biopsy after a standard diagnostic UV provocation using a solar simulator at 1.5 x minimal erythema dose on three consecutive days. Following UV provocation, we observed a striking diffuse increase in the expression of *IFNK* in the epidermis using *in situ* hybridization (**Figure 8B**), similar to the expression observed in non-lesional skin of At-Risk individuals.

**Figure 8.**
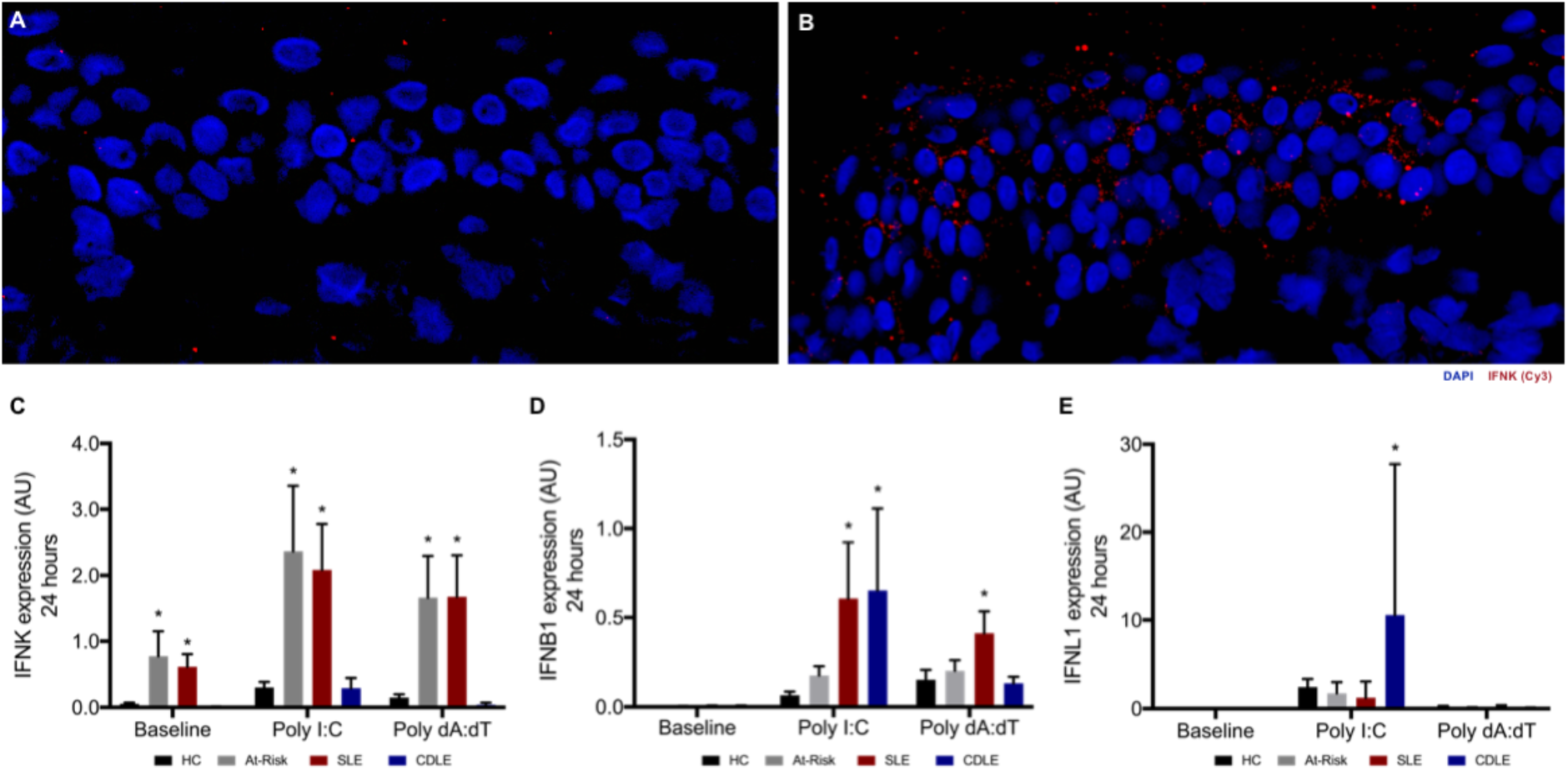
Keratinocytes from At-Risk individuals and SLE patients show high expression of type I IFNs in response to UV light and nucleic acids. (**A**) *IFNK* expression in epidermis of SLE patient with inactive disease before UV provocation. (**B**) *IFNK* expression in epidermis of the same SLE patient after UV provocation. (**C-E**) Human keratinocytes were isolated from fresh skin biopsies and were then cultured in the absence or presence of Poly I:C (1 μg/mL) or Poly dA:dT (100 ng/mL). Expression level of *IFNK* (**C**), *IFNB1* (**D**), *IFNL1* (**E**) in keratinocytes from healthy controls (HC), At-Risk individuals (At-Risk), SLE patients (SLE), and patients with cutaneous discoid lupus erythematosus (CDLE) after *in vitro* culture for 24 hours. Data are represented as mean ± SEM. \**P* < 0.05. 2-way ANOVA (**C-E**).

In order to confirm this cellular source of IFN, we studied the IFN-producing capacity of keratinocytes in At-Risk individuals and SLE patients *in vitro*. We isolated human keratinocytes from non-lesional skin of healthy controls (n = 3), At-Risk individuals (n = 5) and SLE patients (n = 5). Keratinocytes isolated from lesional skin biopsies of patients with cutaneous discoid lupus erythematosus (CDLE), who were ANA-negative and had minimal IFN Score A expression in blood, were also used as a disease control (n = 3). Cells were cultured and stimulated with TLR3 or RIG-I agonists, Poly(I:C) (1 μg/mL) or Poly(dA:dT) (100 ng/ml) respectively, for 6 and 24 hours before the expression of 3 subtypes of type I IFNs (*IFNK*, *IFNA2 and IFNB1*) as well as type III IFN (*IFNL1*) was measured by qRT-PCR.

At baseline, without exogenous stimulation, *IFNK* was expressed by keratinocytes from At-Risk and SLE skin, but not from healthy controls or CDLE. After either Poly(I:C) or Poly(dA:dT) stimulation, this expression of *IFNK* by At-Risk and SLE keratinocytes was further increased (**Figure 8C**). For *IFNB1*, there was no expression at baseline in any sample. However, after stimulation with Poly(I:C) there was a trend to increased expression for At-Risk keratinocytes and a significant increase for keratinocytes from SLE and CDLE patients. *IFNB1* expression was also increased in keratinocytes of SLE patients after Poly(dA:dT) stimulation but not in other conditions (**Figure 8D**). In contrast, *IFNL1* expression was only observed in CDLE keratinocytes following Poly(I:C) stimulation but not in the other conditions or following Poly(dA:dT) stimulation (**Figure 8E**). Finally, *IFNA2* expression by keratinocytes was not found in any sample or condition (data not shown).

## DISCUSSION

The importance of type I IFNs in the pathogenesis of human autoimmune connective tissue diseases such as SLE is now generally accepted based on genetic and gene expression data. But IFNs form a complex system with multiple ligands and receptors with overlapping functions, and all cell types potentially producing and responding to interferons. This complexity has left many unanswered questions about their cellular source, mechanism of dysregulation and role in disease initiation and perpetuation. In this study, by including large cohorts of At-Risk individuals with novel functional, transcriptomic and tissue data, we have been able to answer some of these questions. Several lines of evidence clearly indicated that, even in At-Risk individuals, pDCs have lost their immunogenic functions, do not correlate with other immunologic or clinical features of disease, and are not active in tissue. However, simultaneously we found that in histologically normal skin keratinocytes produced type I IFNs with a marked concentration of IFN response at this site in the absence of infiltrating leucocytes. Our results therefore locate the IFN response in non-haematopoietic tissues prior to the onset of inflammation.

Plasmablasts, other B cell subsets and follicular helper-like T cells are expanded in patients with active SLE, upregulating chemokine receptors and correlating with disease activity (46–49). In contrast, we reported a marked reduction of circulating pDC numbers even at preclinical stage. Previous literature speculated that this may indicate migration of pDCs to inflamed tissues (50, 51). This suggestion was based on the observation of CD123 or CD303 positive cells on immunohistochemistry (52). However, these markers are not specific for pDCs, as we demonstrated that differentiated monocytes express these markers. Moreover, surface markers do not indicate whether these cells are functional. In any case, we found that pDC numbers were already universally low at a preclinical At-Risk stage, with no tissue inflammation and often no long-term progression to clinical autoimmunity. pDCs in both At-Risk and established disease had lost all immunogenic functions with a transcriptomic profile of senescence. These pDCs did not exhibit any change in chemokine receptor expression, and their numbers and function bore no relationship with blood IFN activity, disease activity or therapy.

Other recent data had cast doubt on the contribution of pDCs to the type I IFN activity seen in SLE. While attomolar concentrations of IFN-α protein was detected in pDCs of the monogenic interferonopathy STING, this was not seen in SLE samples. However, responses to TLR agonists were not tested (53). In TREX1-deficient mice there is failure to regulate STING-mediated antiviral response leading to aberrant type I IFN production. This has been shown to initiate in non-haematopoietic cells, similarly to our findings in humans (54). Moreover, experimental work on lupus-prone mice reported a gradual loss of pDC capacity to produce IFN-α at late stage of disease course (55, 56). Importantly, murine models of chronic viral infection maintained a pool of functionally exhausted pDCs – a similar state to what we describe in chronic autoimmunity in humans (57).

Recent findings on Systemic Sclerosis reported the abnormal expression of TLR8 in pDCs that leads to IFN-α production suggesting a key pathological role of RNA-sensing TLR involvement in the establishment of fibrosis (58). However, in our RNA-sequencing data in pDCs sorted from At-Risk individuals or SLE, we could not confirm positive expression of TLR8 in any of the samples.

Within the pDC population, distinct subsets have been described mediating different immune functions (59). Single-cell RNA-sequencing data revealed the diversification of human pDCs in response to influenza virus into three phenotypes (P1-, P2-, P3-pDCs) with distinct transcriptional profiles and functions (60). In our study, pDCs from SLE patients were mostly similar to the P1-phenotype, which represented the conventional secretory function and morphology known about pDCs. This is consistent with our finding that SLE pDCs demonstrated decreased ability to induce CD4^+^CD25^high^FoxP3^+^ T cells, the numbers and function of which are known to be impaired in patients with active SLE (61–63). Additionally, SLE pDCs differentially expressed genes that are well-known to be involved in cellular senescence and stress, negative regulation of TLR and MAPK pathways as well as IL-10 signalling downstream, which can inhibit cytokine production and survival of pDCs (64–66). Oxidative stress is an important feature of lupus pathology, especially in T cells, contributing to a range of biological processes (67). Notably, age-induced cellular stress was shown to affect IFN-α-producing capacity of human pDCs by impairing IRF7 and PI3K pathways (68–70). Similar to our findings in pDCs, kidney infiltrating T cells in murine lupus have recently been shown to exhibit an exhausted transcriptional signature and phenotype (71).

IFN-κ is predominantly produced by human keratinocytes with pleiotropic effects similar to IFN-α/β (72). In SLE patients, keratinocytes have been implicated in the pathogenesis of skin injury by undergoing apoptosis or necrosis and eventually releasing autoantigens (73). Previous studies demonstrated that keratinocytes from patients with cutaneous lupus erythematosus presented increased production of IL-6 compared to healthy controls, with type I IFNs enhancing this process (74). In addition, *IFNK* expression was reported to be significantly increased in lesional skin of patients with cutaneous lupus erythematosus related to photosensitivity (75). By visualising the direct expression of type I IFN transcripts in human skin biopsies, we confirmed and expanded these findings. Diffuse expression of *IFNK* was seen not only in epidermis of lesional skin of SLE patients but more importantly, in epidermis of non-lesional skin of ANA-positive At-Risk individuals with high IFN activity in blood, who had no clinical or histopathological features of inflammation. We demonstrated that UV provocation *in vivo* induced higher expression of *IFNK* in epidermis of SLE patients with inactive disease. Notably, *in vitro* culture of keratinocytes showed high expression of *IFNK* at baseline, whilst stimulation with TLR3 and RIG-I agonists enhanced IFNK expression even in At-Risk individuals. Further, the types of IFN produced varied between systemic and discoid lupus and stimuli. These results therefore indicate production of IFN by non-haematopoietic cells in the absence of production by pDCs or tissue leucocytes early in the initiation of autoimmunity and in a disease-specific manner.

In summary, while the importance of type I IFN in SLE is undeniable, the reasons for the failure of normal regulation of its production have never been clear. Our results indicate that tissues such as the skin are not mere passive targets for leucocyte mediated immune processes but play an active role in generating an IFN response which dominates over inert pDCs. Immune functions of non-haematopoietic tissues may play a role in determining the clinical presentation of these heterogeneous diseases. They may help to explain the resistance of tissue inflammation to commonly used therapies that target leucocytes, and instead point to novel therapeutic targets that lie outside the conventional immune system. Further study of these tissues may offer new insights into the early events in the initiation of autoimmune disease and therefore how these might be targeted for disease prevention.

## METHODS

### Patients and controls

Peripheral blood and skin biopsies were obtained from healthy individuals and patients from different disease groups (SLE, pSS, At-Risk). Patients were recruited based on 2012 SLICC classification criteria for SLE, 2016 ACR/EULAR classification criteria for pSS, while At-Risk individuals were classified as ANA positive, ≤1 SLE clinical criterion, symptom duration <12 months and treatment-naïve. **Supplemental Table 1** summarizes the characteristics and treatment of SLE patients. Ethical approval was provided by Leeds East – National Research Ethics Committee (REC 10/H1306/88).

### Isolation of human peripheral blood cells

Human PBMCs were separated from whole blood by a density gradient centrifugation method using Leucosep tubes (Greiner Bio-One). pDCs were purified from freshly isolated PBMCs by negative selection using the Diamond Plasmacytoid Dendritic Cell Isolation Kit II (Miltenyi Biotec). Naïve CD4^+^ T cells were purified by negative selection using the Naive CD4^+^ T Cell Isolation Kit II (Miltenyi Biotec). Pre-enriched pDCs were sorted using an antibody to BDCA-4 (Miltenyi Biotec). Cell sorting was carried out at the SCIF Flow Cytometry and Imaging Facility of the Wellcome Trust Brenner Building, University of Leeds, with a BD Influx 6 Way Cell Sorter (BD Biosciences).

### Culture of human peripheral blood cells

Cells were cultured in RPMI medium 1640 with GlutaMAX supplement (ThermoFisher Scientific) containing 10% (vol/vol) FBS and 100 U/ml penicillin/streptomycin. For cytokine production, PBMCs were stimulated with 2 μM class A CpG (ODN 2216; Miltenyi Biotec) or 2 μM ORN R-2336 (Miltenyi Biotec). For pDC/T-cell co-culture, purified pDCs (1 × 10^5^) were cultured with autologous or allogeneic naïve CD4^+^ T cells (5 × 10^5^) for 5 days in the absence or presence of anti-CD3/CD28 beads (T cell activation/expansion kit; Miltenyi Biotec) at a bead-to-cell ratio of 1:2. Cytokine production was measured by intracellular staining.

### Flow cytometry

For cell surface staining, fluorochrome-conjugated monoclonal antibodies against human CD3, CD4, CD19, CD14, CD56, CD11c, HLA-DR, CD123, CD303, CD304, CD85g, CD85j, CD69, CD25 (Miltenyi Biotec), CD317 (BioLegend) and isotype controls were used. For intracellular staining, cells were first stained for surface markers and then fixed and permeabilized using the Intracellular Fixation & Permeabilization Buffer Set (eBioscience). Fluorochrome-conjugated monoclonal antibodies against human IFN-α, TNF, IL-6, IFN-γ, IL-10 (Miltenyi Biotec), TLR9 (BD Biosciences), TLR7 (R&D Systems) and isotype controls were used. For FoxP3 (Miltenyi Biotec) intracellular staining, cells were first stained for surface markers and then fixed and permeabilized using the FoxP3 Staining Buffer Set (Miltenyi Biotec). Cell proliferation was measured using the CellTrace Violet Cell Proliferation kit (ThermoFisher Scientific) according to the manufacturer’s instructions. Flow cytometry was performed on LSRII (BD Biosciences) or Cytoflex S (Beckman Coulter) and the data were analysed using FACS DiVA (BD Biosciences) or CytExpert (Beckman Coulter) software.

### Quantification of gene expression in peripheral blood

Total RNA was extracted from freshly isolated PBMCs using the Total RNA Purification Kit (Norgen Biotek) according to manufacturer’s instructions. RNA was reverse-transcribed to cDNA using the Fluidigm Reverse Transcription Master Mix buffer and gene expression was measured by TaqMan assays (Applied Biosystems, Invitrogen). The relative expression of specific transcripts (*ISG15*, *IFI44*, *IFI27*, *CXCL10*, *RSAD2*, *IFIT1*, *IFI44L*, *CCL8*, *XAF1*, *GBP1*, *IRF7*, *CEACAM1*) was normalized with respect to the internal standard (*PP1A*). IFN Score A was calculated based on Factor Analysis previously described (8).

### RNA-sequencing data generation

RNA from sorted pDCs was extracted using PicoPure RNA Isolation Kit (ThermoFisher Scientific) and quantified using Qubit RNA HS Assay Kit (Thermo Fisher Scientific). RNA libraries were made by using SMART-Seq V4 ultra low Input RNA Kit (Takara Bio USA) and Nextera XT DNA Library Preparation Kit (Illumina) for NGS sequencing. Indexed sequencing libraries were pooled and sequenced on a single lane on HiSeq 3000 instrument as 151bp paired-end reads. Pooled sequence data was then demultiplexed using Illumina bcl2fastq software allowing no mismatches in the read index sequences.

### Raw-sequencing data processing and analysis

Raw paired-end sequence data in Fastq format was initially analysed using FastQC software in order to identify potential issues with data quality. Cutadapt software was then used to remove poor quality bases (Phred quality score <20) and contaminating technical sequences from raw sequenced reads. Contaminating technical sequences identified at the initial QC stage were as follows:

CTGTCTCTTATA – Next Era Transposase Sequence

GTATCAACGCAGAGTACT– SmartSeq Oligonucleotide Sequence

dT30 – SmartSeq 3’ CDS Primer II sequence

Reads trimmed to fewer than 30 nucleotides and orphaned mate-pair reads were discarded to minimise alignment errors downstream.

Reads were aligned to human hg38 analysis set reference sequences, obtained from UCSC database (76) using splicing-aware STAR aligner (77) for RNA-Sequencing data. STAR aligner was run in 2-pass mode, with known splice junctions supplied in GTF file format, obtained from hg38 RefSeq gene annotation table from UCSC database using Table Browser tool (78). The resulting alignments in BAM file format were checked for quality using QualiMap software (79) and Picard tools (80). Picard tools were used to mark PCR/Optical duplicate alignments. Custom code was used to filter out contaminating ribosomal RNA alignments, using ribosomal RNA coordinates for hg38 analysis set reference obtained using UCSC Table Browser tool. The final alignment files were sorted and indexed using Samtools software (81) and visualised using IGV browser (82).

Bioconductor R package RSubread (83) was used to extract raw sequenced fragment counts per transcript using RefSeq hg38 transcript annotation set, as before. Paired-end reads were counted as a single fragment and multi-mapping read pairs were counted as a fraction of all equivalent alignments. Raw count data was normalised for library size differences using median ratio method (84), as implemented in DESeq2 R Bioconductor package (85). DESeq2 was also used to perform additional data QC steps and differential expression analyses. Differentially expressed gene expression was visualised as clustered heatmaps using Pheatmap R package (86) using log-transformed normalised gene expression values as input. Gene functional and pathway enrichment analyses were performed using R Bioconductor packages clusterProfiler (87) and ReactomePA (88). Additionally, KEGG (89) pathways were visualised using Pathview package (90).

### Measurement of relative telomere length

Relative telomere length was measured using Telomere PNA Kit/FITC for Flow Cytometry (Agilent) according to manufacturer’s protocol. Briefly, on a single cell suspension consisting of purified pDCs and control cells (1301 cell line; Sigma-Aldrich), the sample DNA was denatured for 10 minutes at 82 °C either in the presence of hybridization solution without probe or in hybridization solution containing fluorescein-conjugated PNA telomere probe. Then hybridization took place in the dark at room temperature overnight. The sample was then resuspended in appropriate buffer for further flow cytometric analysis. The data obtained were used for determination of the relative telomere length as the ratio between the telomere signal of each sample (pDCs) and the control cell (1301 cell line) with correction for the DNA index of G_0/1_ cells.

### Oxidative stress assay

Freshly isolated PBMCs from healthy donors were exposed to H_2_O_2_ (0 − 500 μM) for 15 minutes. After exposure, cells were washed thoroughly and resuspended at 1 × 10^6^ in culture medium before they were stimulated with 2μM ODN 2216 (Miltenyi Biotech) for 6 hours. The production of IFN-α by pDCs was measured by intracellular staining as described above.

### UV provocation

UV provocation was performed based on a published protocol designed for use in clinical trials (91, 92). Briefly, a solar simulator was used in routine clinical practice, which replicated the protocol of UV-A and UV-B provocation in a single exposure. On day 1, four 1.5 cm^2^ areas of skin were exposed to solar simulated radiation depending on skin type; 4, 8, 12, 16 J/cm^2^ for skin types I and II, and 6, 12, 18, 24 J/cm^2^ for skin types III-VI. On day 2, the minimal erythema dose was then determined. A 10 cm^2^ non-sun exposed area of skin was exposed to minimal erythema dose × 1.5 on three consecutive days. A biopsy of the pre-exposed and exposed area of skin was obtained when a reaction was seen clinically (mean time to a positive reaction to provocation was 7 (±6) days, and rarely more than 14 days).

### Tissue section

Skin biopsies were obtained from healthy individuals and patients, then snap frozen in liquid nitrogen within 5 minutes, embedded in OCT and stored in −80°C freezer. Fresh frozen skin biopsies were cryosectioned to 10–20 μΜ, placed on superfrost plus slides (Thermo Scientific) and used for *in situ* hybridization.

### *In situ* hybridization and fluorescence microscopy

*In situ* hybridization of type I IFNs transcripts in skin samples was performed using RNAscope Multiplex Fluorescent Reagent Kit v2 (Advanced Cell Diagnostics) according to manufacturer’s instructions. Briefly, cryosections were fixed and dehydrated before exposure to hydrogen peroxide and protease treatment followed by hybridization for 2 hours with custom-designed target probes (*IFNA2*, *IFNK*). Appropriate positive and negative controls were provided by the manufacturer. Hybridization signals were amplified and detected using TSA Plus fluorescein and TSA Plus Cyanine 3 (Perkin Elmer) according to manufacturer’s protocol. Nuclei were highlighted using 4’,6-diamidino-2-phenylindole (DAPI). Slides were mounted using Prolong Gold Antifade mounting medium (ThermoFischer Scientific) and dried overnight in the dark at 4°C. Images were acquired on a Nikon A1R confocal laser scanning microscope system at 20–40x magnification. Images were analysed in Nikon NIS Elements software.

### Culture of human keratinocytes

Human keratinocytes were isolated from 3 mm punch skin biopsies. For keratinocytes, the epidermal component of the biopsy was placed in a T75 flask and cultured at 37°C in low glucose DMEM (Fischer Scientific) containing 10% (vol/vol) FBS (Fischer Scientific) and 1% penicillin/streptomycin. Keratinocytes were passaged and sub-cultured into keratinocyte growth medium (PromoCell) for continuous culture. Keratinocytes were passaged and plated in 24-well plates for subsequent stimulation. At 90% confluence, cells were either untreated or treated with 1 μg/ml Poly I:C (InvivoGen) or 100 ng/ml Poly dA:dT (InvivoGen) for 6 or 24 hours.

### Quantitative RT-PCR for keratinocytes

RNA was extracted from keratinocytes using Quick-RNA MiniPrep kit (Zymo Research) according to manufacturer’s instructions. Extracted RNA was reverse transcribed using First Strand cDNA Synthesis kit (ThermoFisher). The cDNA was then used in qRT-PCR assay using QuantiFast SYBR Green PCR kit (Qiagen). For the assay, the following QuantiTech primers were used: *IFNK* (QT00197512; Qiagen), *IFNB1* (QT00203763; Qiagen), *IFNL1* (QT00222495; Qiagen), *IFNA2* (QT00212527; Qiagen), *U6snRNA* (forward—5′-CTCGCTTCGGCAGCACA-3′; reverse—5′-AACGCTTCACGAATTTGC-3′; Sigma-Aldrich). For gene expression analysis, ddCt method was used and all samples were normalised to the housekeeping gene (*U6snRNA*).

### Statistical analysis

Statistical analyses were carried out with Prism software (GraphPad). Continuous variables were compared using either Student’s T test or ANOVA followed by pairwise Tukey tests. Pearson’s correlation was used for associations. A p value of ≤ 0.05 was considered significant (ns, not significant; \**P* < 0.05; \*\**P* < 0.01; \*\*\**P* < 0.001; \*\*\*\**P* < 0.0001). In all Fig., error bars indicate SEM.

## AUTHOR CONTRIBUTIONS

Dr Psarras planned and performed most of the *in vitro* experiments, conducted data analysis and wrote the manuscript. Dr Alase performed *in vitro* experiments. Dr Antanaviciute and Dr Carr performed RNA-sequencing and data analysis. Dr Md Yusof and Dr Vital provided patient samples and clinical data. Dr Wittmann, Prof Emery, Prof Tsokos planned experiments, provided scientific input and wrote the manuscript. Dr Vital supervized the project, planned experiments, conducted data analysis and wrote the manuscript. All authors read and approved the manuscript.

## ACKNOWLEDGEMENTS

Cell sorting was performed by SCIF Flow Cytometry Facility of the Wellcome Trust Brenner Building at the University of Leeds by Dr Adam Davison and Ms Elizabeth Straszynski. Dr Yasser El-Sherbiny provided flow cytometry support. Ms Ummey Hany provided technical support for RNA-sequencing. Dr Vital was funded by an NIHR Clinician Scientist Fellowship (CS-2013–13–032) and Dr Yusof was funded by an NIHR Doctoral Research Fellowship (DRF-2014–07–155). The views expressed are those of the author(s) and not necessarily those of the NHS, the NIHR or the Department of Health.

