## Supplemental Figures and Tables for "Plasmacytoid dendritic cells are functionally exhausted while non-haematopoietic sources of type I interferon dominate human autoimmunity"

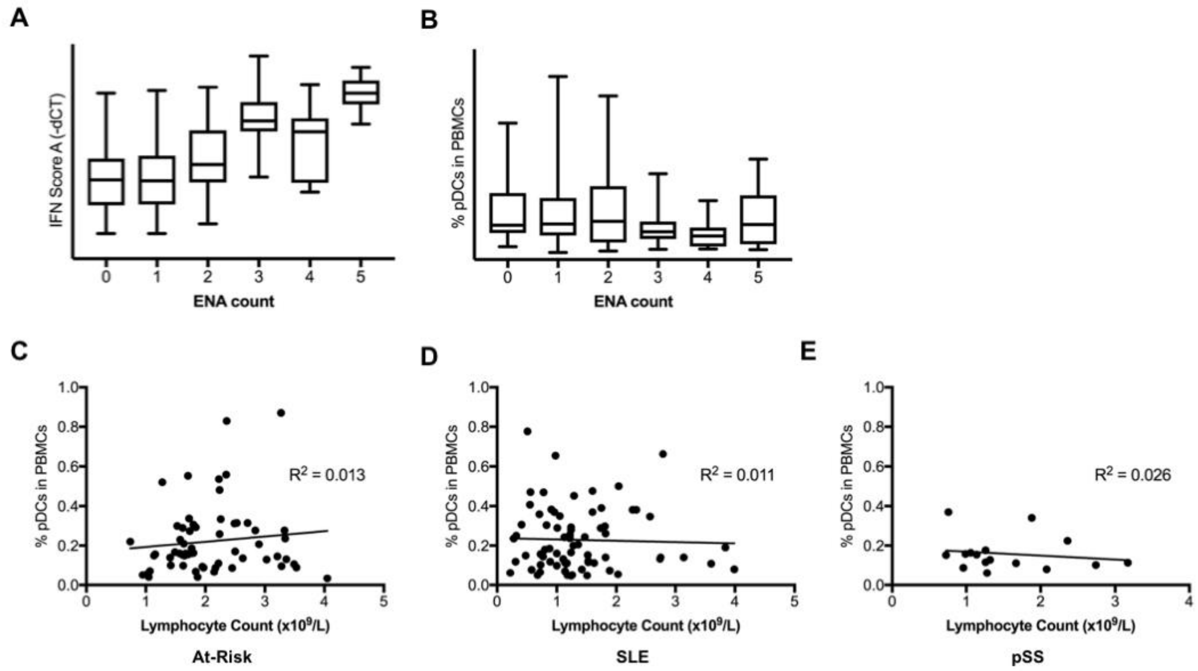


**Supplemental Figure 1.** Association of peripheral pDCs with other immunological parameters. (**A**) Higher IFN Score A was correlated to a higher ENA count. (**B**) The percentage of pDCs in peripheral blood showed no correlation with ENA count. (**C**) Association of percentage of pDCs in peripheral blood and lymphocyte count in At-Risk individuals. (**D**) Association of percentage of pDCs in peripheral blood and lymphocyte count in patients with SLE. (**E**) Association of percentage of pDCs in peripheral blood and lymphocyte count in patients with pSS. Data are represented as mean ± SEM. Nonlinear regression (**C-E**).

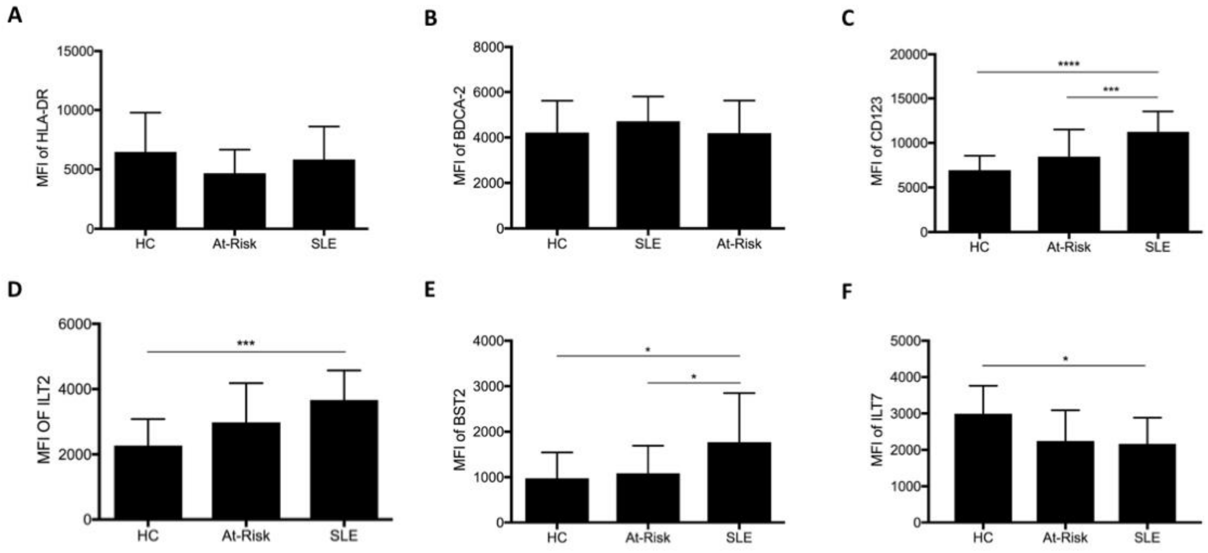


**Supplemental Figure 2.** Phenotyping peripheral pDCs in At-Risk individuals and patients with SLE in comparison with healthy controls (HC) for the expression of HLA-DR (**A**), BDCA-2 (**B**), CD123 (**C**), ILT2 (**D**), BST2 (**E**), and ILT7 (**F**). Data are represented as mean ± SEM. *$P$ < 0.05; ***$P$ <0.001. 2-way ANOVA (**A-F**).

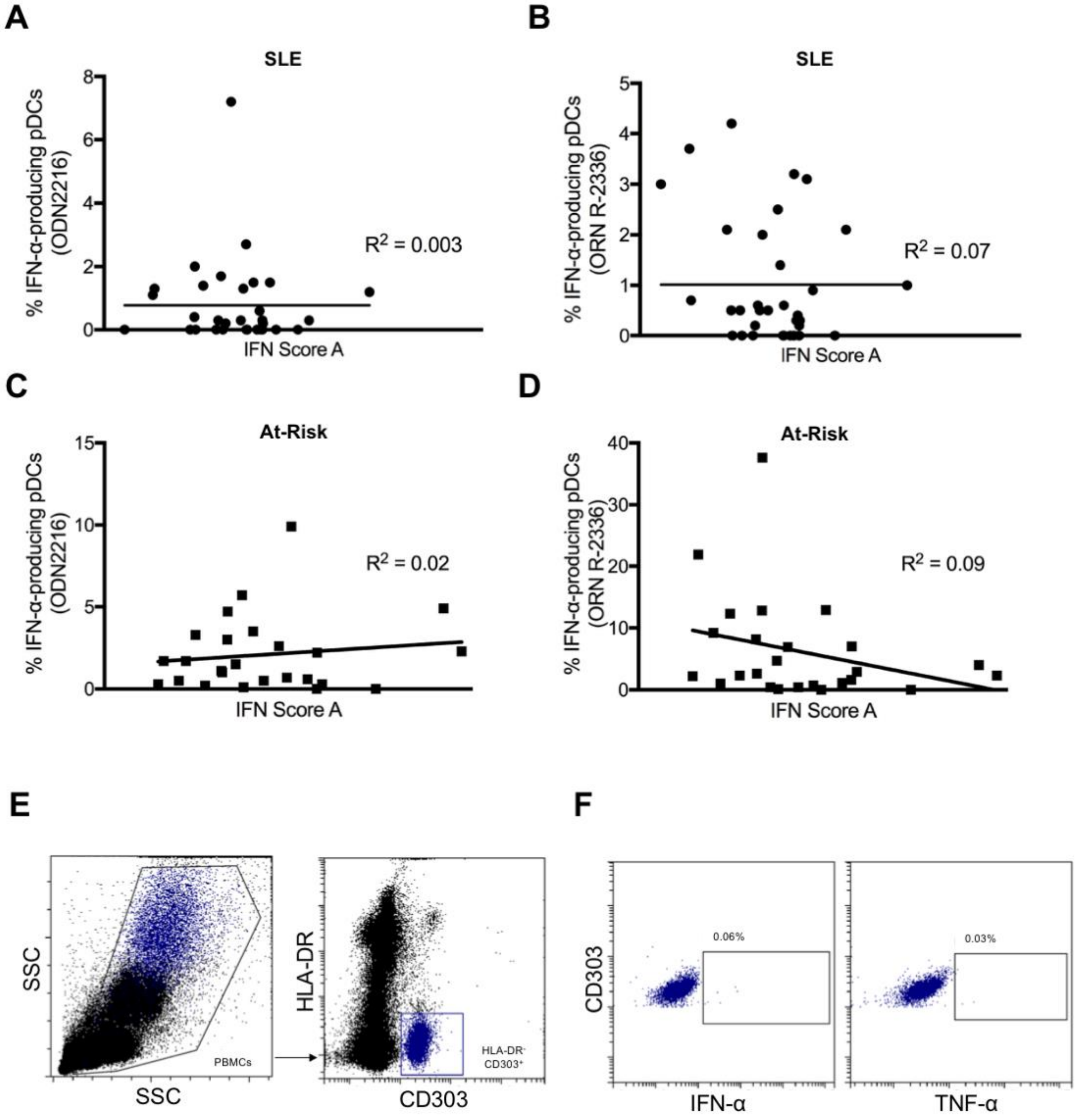


**Supplemental Figure 3.** (**A-D**) No association between TLR-mediated IFN-α production and IFN Score A in SLE patients and at-risk individuals. (**E**) Gating of HLA-DR$^-$CD303$^+$ cells from cultured PBMCs. (**F**) No production of IFN-α or TNF-α was detected by HLA-DR$^-$CD303$^+$ cells after TLR9 stimulation (ODN 2216). Data are represented as mean ± SEM. Nonlinear regression (**A-D**).

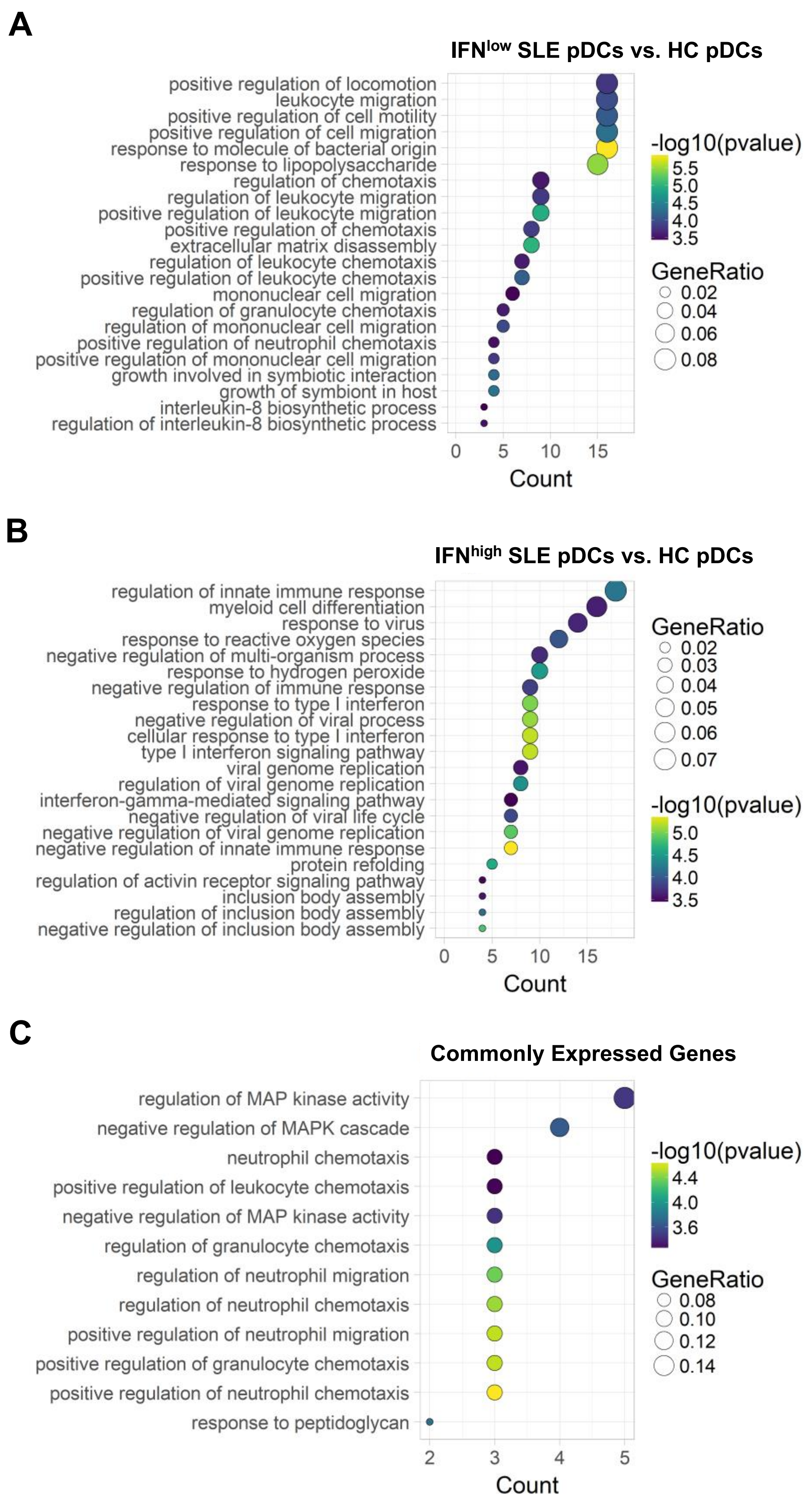


**Supplemental Figure 4.** Gene Ontology Biological Process Term Enrichment in differentially expressed genes in pDCs of IFN$^{low}$ SLE patients (**A**), IFN$^{high}$ SLE patients (**B**) and genes commonly expressed to both (**C**) in comparison with pDCs from healthy controls.

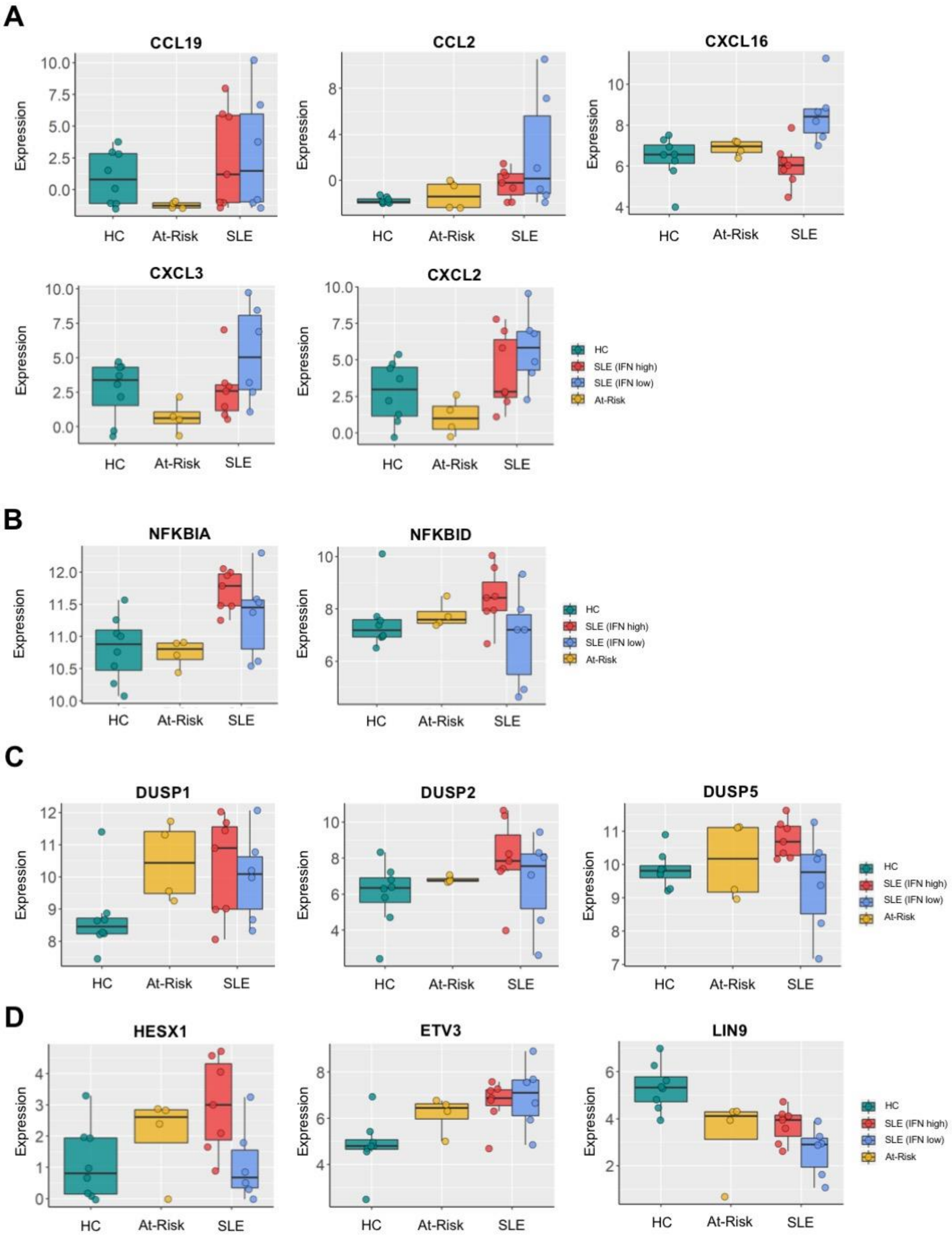


**Supplemental Figure 5.** Differentially expressed genes in pDCs of healthy controls (HC), at-risk individuals (At-Risk), IFN$^{low}$ SLE and IFN$^{high}$ SLE patients: (**A**) Chemokines; (**B**) NF-κB inhibitors; (**C**) Phosphatases; (**D**) Transcriptional repressors.

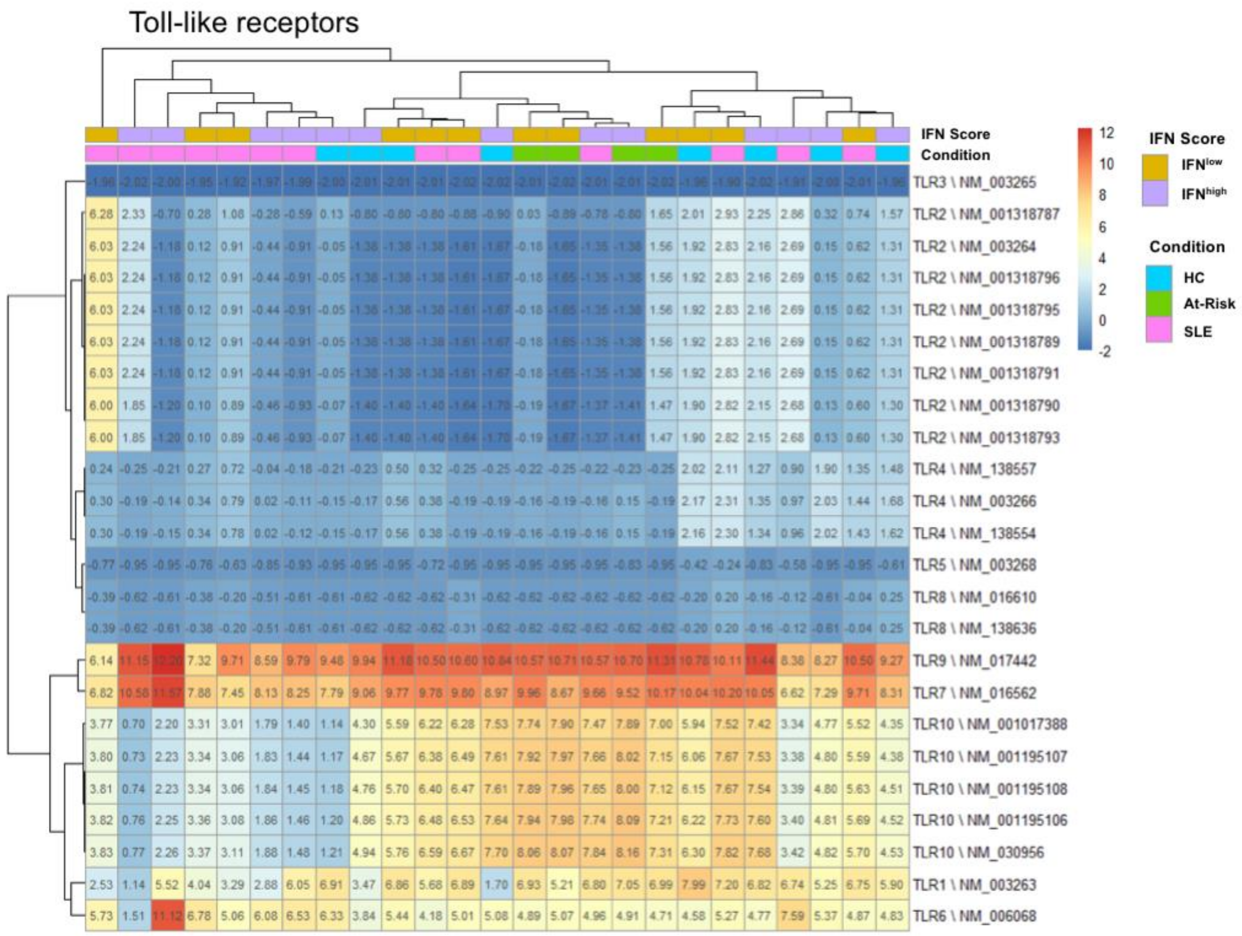


**Supplemental Figure 6.** Differential expression of Toll-like receptors (TLRs) in pDCs of healthy controls (HC), at-risk individuals (At-Risk), and SLE patients.

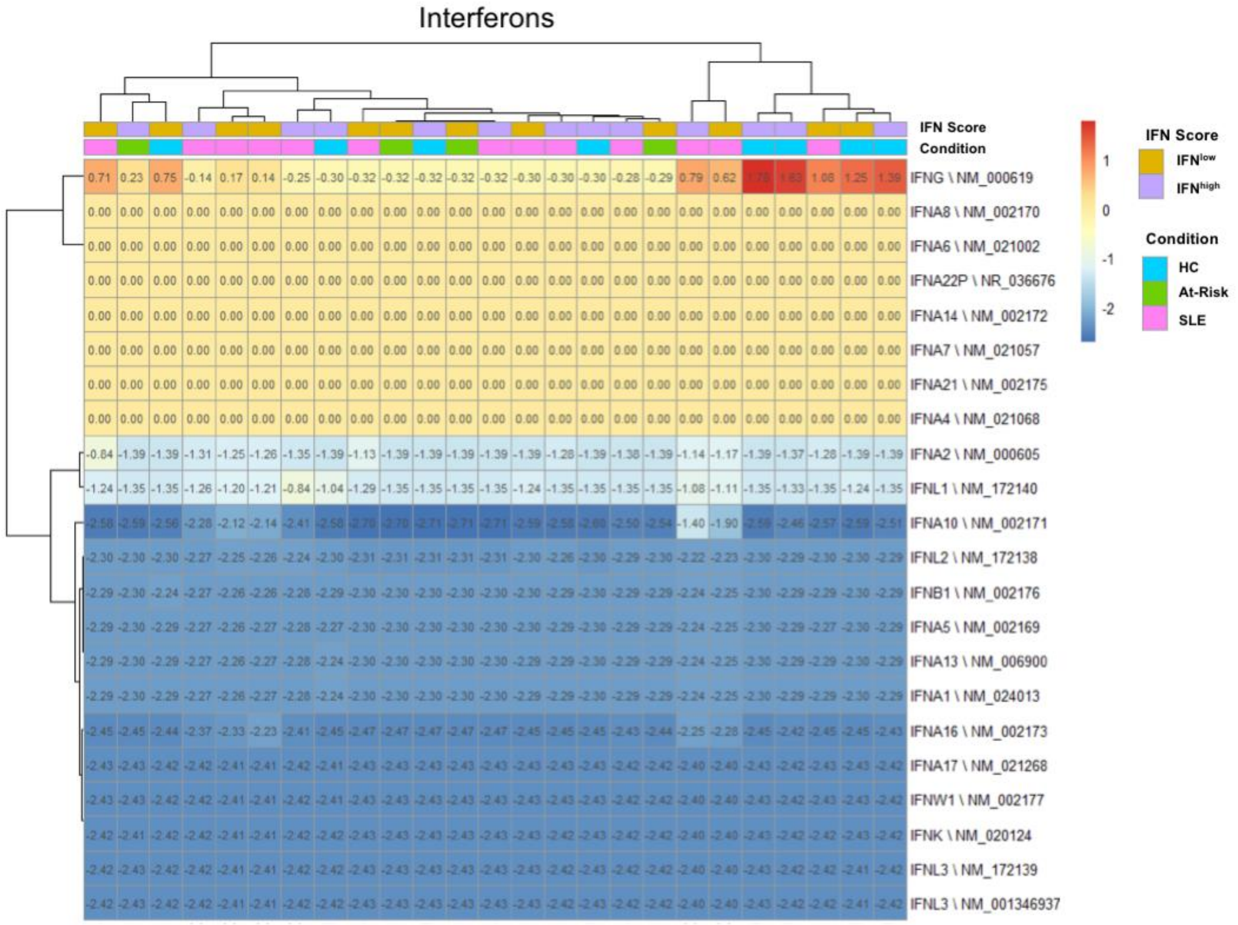


**Supplemental Figure 7.** Differential expression of type I, II, and III IFNs in pDCs of healthy controls (HC), at-risk individuals (At-Risk), and SLE patients.

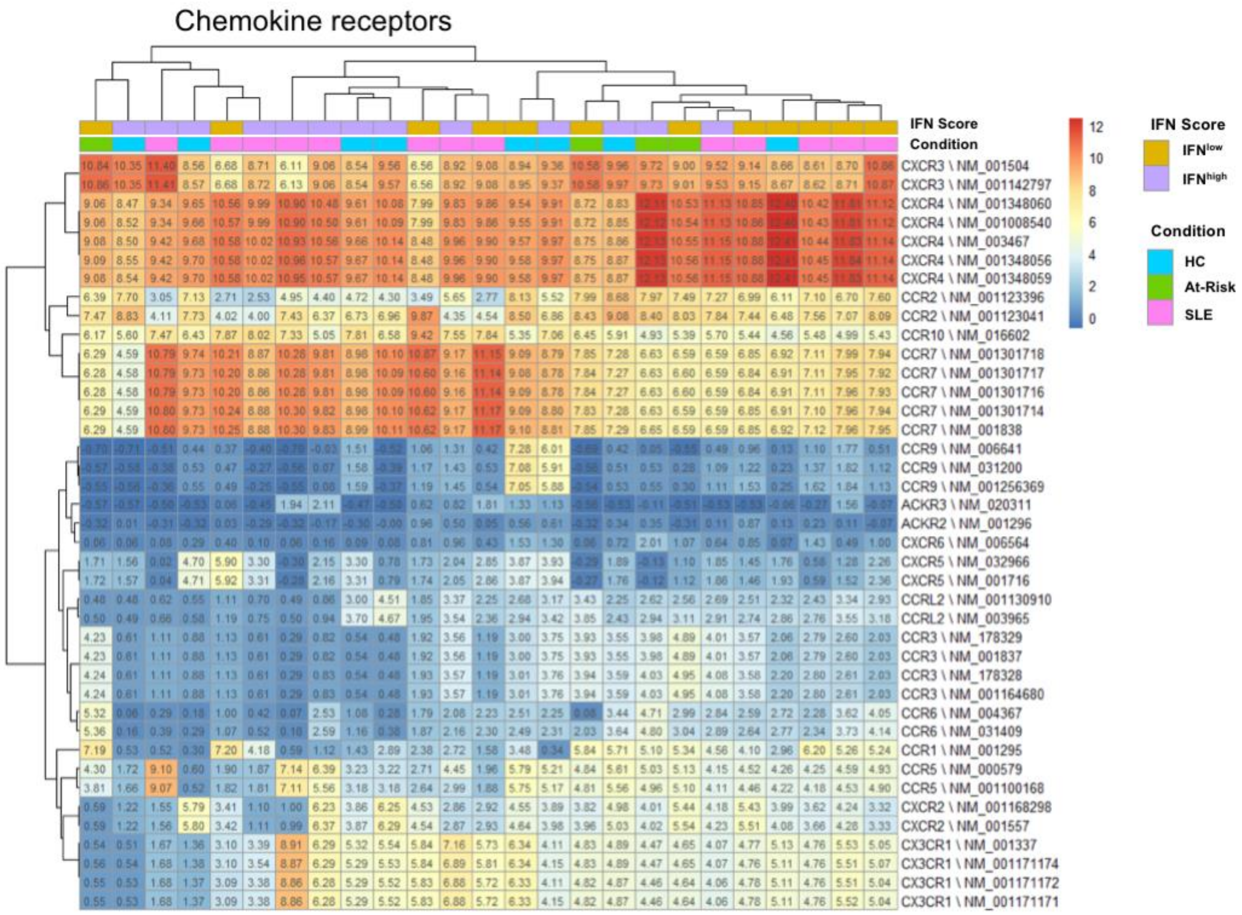


**Supplemental Figure 8.** Differential expression of chemokine receptors in pDCs of healthy controls (HC), at-risk individuals (At-Risk), and SLE patients.

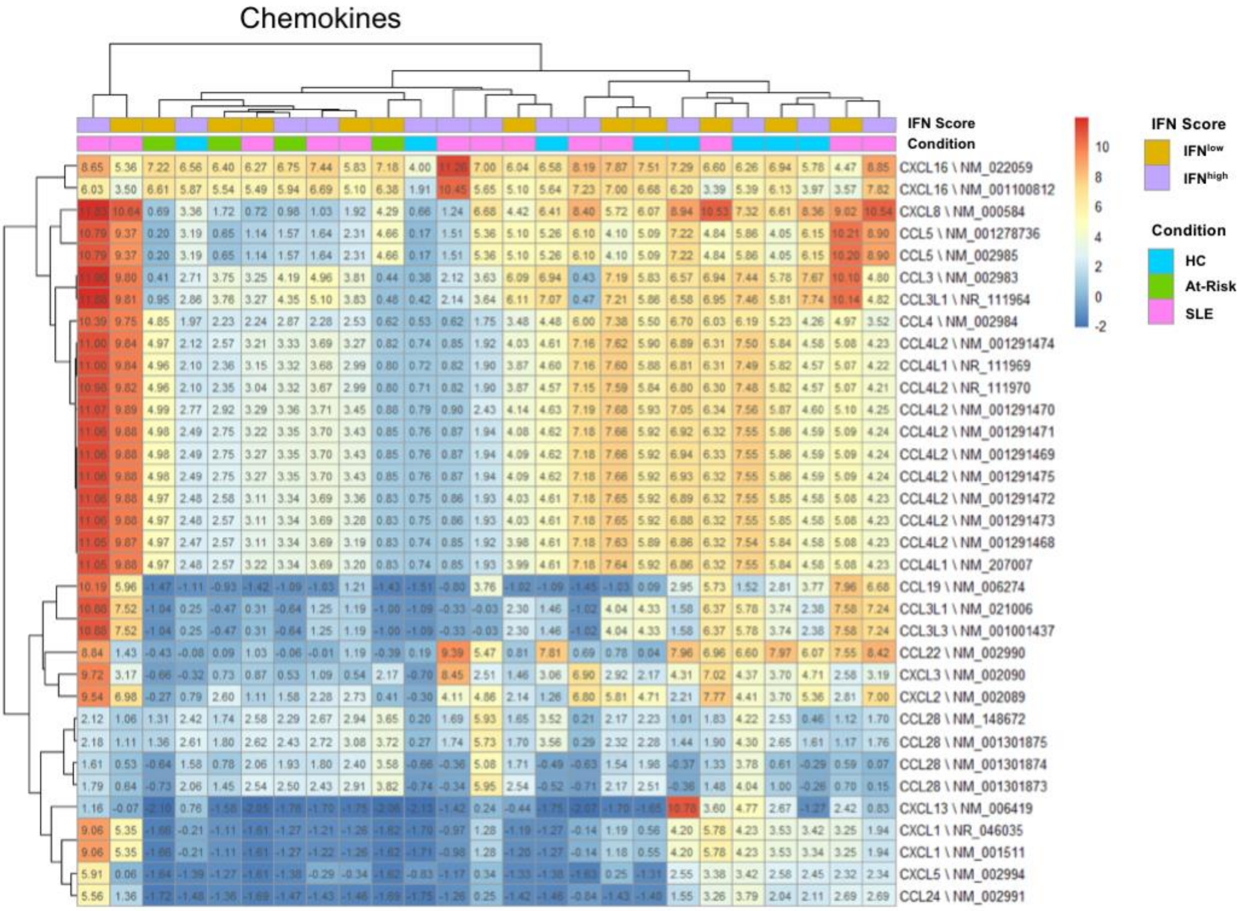


**Supplemental Figure 9.** Differential expression of chemokines in pDCs of healthy controls (HC), at-risk individuals (At-Risk), and SLE patients.

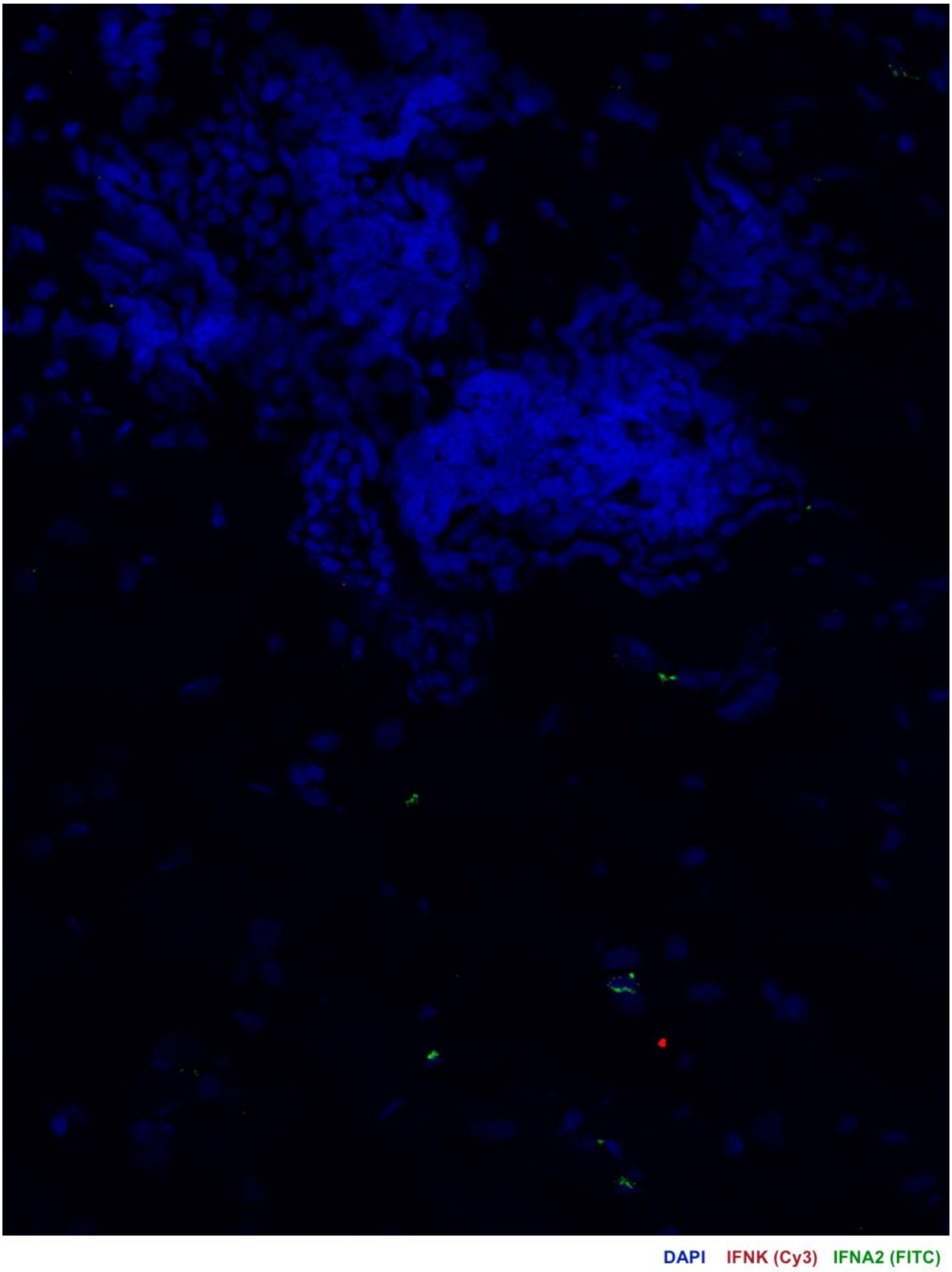


**Supplemental Figure 10.** Area of leucocyte infiltration and connective tissue of a patient with SLE with active skin lesion. Skin biopsies were hybridized using RNAscope *in situ* hybridization technology with custom-designed target probes for *IFNA2* and *IFNK*. Hybridization signals were amplified and detected using TSA Plus fluorescein (FITC) for IFNA2 and TSA Plus Cyanine 3 (Cy3) for *IFNK*. Nuclei were highlighted using DAPI. *IFNA2* expression was detected in cells within the connective tissue but not in infiltrating leucocytes.

| | |
|---|---|
| Age, median (range) years | 44 (18 -76) |
| Female patients (%) | 94 |
| Ethnicity (%) | |
| Caucasian | 74 |
| South Asian | 16 |
| East Asian | 5 |
| African/Caribbean | 5 |
| Clinical symptoms: BILAG Score A/B (%) | |
| Mucocutaneous | 23 |
| Musculoskeletal | 23 |
| Haematological | 3 |
| Renal | 16 |
| Neurological | 3 |
| Cardiorespiratory | 2 |
| Gastrointestinal | 3 |
| Opthalmic | 0 |
| General | 0 |
| Hydroxychloroquine (%) | 83 |
| Other immunosuppressants (%) | |
| Methotrexate | 14 |
| Azathioprine | 18 |
| Mycophenolate mofetil | 27 |
| Cyclophosphamide | 1 |
| Oral steroids (%) | 47 |

**Supplemental Table 1.** Clinical characteristics and treatment of SLE patients.

| | IFN$^{low}$ SLE pDCs vs. HC pDCs | | |
|---|---|---|---|
| **Gene** | **Fold Change (log2)** | ***P* value** | **FDR** |
| ULBP2 | 19.757 | < 0.001 | < 0.001 |
| C10orf35 | 18.302 | < 0.001 | < 0.001 |
| TPSB2 | 18.228 | < 0.001 | < 0.001 |
| TMEM216 | -4.981 | < 0.001 | < 0.001 |
| CXCL2 | 6.379 | < 0.001 | < 0.001 |
| PLA2G7 | 9.712 | < 0.001 | < 0.001 |
| DNAJB4 | 1.965 | < 0.001 | 0.001 |
| ZBP1 | -7.254 | < 0.001 | 0.001 |
| LUCAT1 | 6.207 | < 0.001 | 0.006 |
| LOC100861532 | 2.839 | < 0.001 | 0.007 |
| LGALSL | 6.161 | < 0.001 | 0.007 |
| SNORD95 | 4.390 | < 0.001 | 0.007 |
| LOC100008587 | 3.428 | < 0.001 | 0.007 |
| BBC3 | 2.962 | < 0.001 | 0.011 |
| SPR | 6.792 | < 0.001 | 0.015 |
| FN1 | 4.352 | < 0.001 | 0.015 |
| PHF3 | 1.125 | < 0.001 | 0.015 |
| HSPB1 | 1.527 | < 0.001 | 0.015 |
| CLEC5A | 8.287 | < 0.001 | 0.015 |
| CKAP4 | -6.563 | < 0.001 | 0.015 |
| IRAIN | 5.493 | < 0.001 | 0.015 |
| FAM157C | 3.945 | < 0.001 | 0.015 |
| DNAJB1 | 2.108 | < 0.001 | 0.015 |
| ETV3 | 3.037 | < 0.001 | 0.017 |
| TPSAB1 | 9.650 | < 0.001 | 0.020 |
| SLC6A6 | 3.758 | < 0.001 | 0.020 |
| NPAS1 | 5.880 | < 0.001 | 0.020 |
| GADD45B | 2.245 | < 0.001 | 0.021 |
| IFITM1 | -3.619 | < 0.001 | 0.022 |
| COCH | -7.844 | < 0.001 | 0.022 |
| ANO8 | 3.516 | < 0.001 | 0.022 |
| LINC00623 | 3.718 | < 0.001 | 0.022 |
| HSPA4 | -2.406 | < 0.001 | 0.023 |
| AOAH | -6.704 | < 0.001 | 0.023 |
| HOXB3 | 4.793 | < 0.001 | 0.023 |
| SLC8A2 | 5.243 | < 0.001 | 0.023 |
| NPL | 4.663 | < 0.001 | 0.025 |
| ZNF678 | 4.063 | < 0.001 | 0.025 |
| CNST | 2.130 | < 0.001 | 0.025 |
| RNY5 | 3.224 | < 0.001 | 0.025 |
| PLEKHA8P1 | -4.947 | < 0.001 | 0.025 |
| DUSP1 | 2.903 | < 0.001 | 0.027 |
| LOC100008589 | 2.695 | < 0.001 | 0.028 |
| NRP2 | 6.969 | < 0.001 | 0.028 |
| EGR2 | 7.175 | < 0.001 | 0.029 |
| FCGR2B | -6.697 | < 0.001 | 0.033 |
| NRG2 | 5.224 | < 0.001 | 0.033 |
| LOC221946 | 5.139 | < 0.001 | 0.033 |
| PLCXD1 | 3.230 | < 0.001 | 0.033 |
| LACC1 | 4.975 | < 0.001 | 0.033 |

**Supplemental Table 2.** Top 50 genes that are differentially expressed in pDCs of IFN$^{low}$ SLE patients in comparison with pDCs from healthy controls (HC).

| | IFN$^{high}$ SLE pDCs vs. HC pDCs | | |
|---|---|---|---|
| **Gene** | **Fold Change (log2)** | ***P* value** | **FDR** |
| IFI44 | 5.403 | < 0.001 | < 0.001 |
| CAMP | 7.261 | < 0.001 | < 0.001 |
| OASL | 6.812 | < 0.001 | < 0.001 |
| SLC8A2 | 5.644 | < 0.001 | < 0.001 |
| ATF3 | 5.870 | < 0.001 | < 0.001 |
| CMPK2 | 7.710 | < 0.001 | < 0.001 |
| BCL2 | 5.485 | < 0.001 | < 0.001 |
| LGALSL | 6.240 | < 0.001 | < 0.001 |
| ORM1 | 8.011 | < 0.001 | < 0.001 |
| NCMAP | 4.931 | < 0.001 | < 0.001 |
| HCG11 | -3.108 | < 0.001 | < 0.001 |
| PARP14 | 3.402 | < 0.001 | < 0.001 |
| MSL2 | 2.238 | < 0.001 | 0.001 |
| DDX60L | 6.470 | < 0.001 | 0.001 |
| PPDPF | 1.798 | < 0.001 | 0.001 |
| IER2 | 2.552 | < 0.001 | 0.001 |
| NR4A1 | 5.230 | < 0.001 | 0.002 |
| TFB1M | -2.213 | < 0.001 | 0.002 |
| GFOD1 | 4.279 | < 0.001 | 0.002 |
| TP53INP2 | 5.236 | < 0.001 | 0.002 |
| TMEM177 | -3.452 | < 0.001 | 0.002 |
| LIPT1 | -2.499 | < 0.001 | 0.002 |
| JUND | 2.486 | < 0.001 | 0.002 |
| THG1L | -2.089 | < 0.001 | 0.002 |
| CCDC121 | 6.073 | < 0.001 | 0.002 |
| SNORD14C | 4.151 | < 0.001 | 0.002 |
| C2orf74 | -2.833 | < 0.001 | 0.002 |
| CISD1 | -2.392 | < 0.001 | 0.002 |
| HIST1H4F | -2.531 | < 0.001 | 0.002 |
| SEMA7A | 3.150 | < 0.001 | 0.002 |
| ETV3L | 3.865 | < 0.001 | 0.002 |
| CEACAM1 | 5.448 | < 0.001 | 0.002 |
| L3MBTL2 | -1.887 | < 0.001 | 0.002 |
| ETV3 | 2.959 | < 0.001 | 0.002 |
| NPAS1 | 5.469 | < 0.001 | 0.002 |
| QRICH2 | 4.336 | < 0.001 | 0.002 |
| LTC4S | 5.089 | < 0.001 | 0.003 |
| ARID5A | 1.781 | < 0.001 | 0.003 |
| EIF2AK2 | 2.358 | < 0.001 | 0.003 |
| HES4 | 4.895 | < 0.001 | 0.003 |
| LINC00847 | -2.487 | < 0.001 | 0.003 |
| NSUN7 | 5.334 | < 0.001 | 0.003 |
| DNAJB1 | 4.229 | < 0.001 | 0.003 |
| KLF8 | -4.556 | < 0.001 | 0.003 |
| RSAD2 | 4.959 | < 0.001 | 0.003 |
| JUN | 2.582 | < 0.001 | 0.003 |
| MON1A | -2.248 | < 0.001 | 0.003 |
| IFI44L | 3.195 | < 0.001 | 0.004 |
| WDR87 | 6.155 | < 0.001 | 0.004 |
| IRAIN | 4.774 | < 0.001 | 0.004 |

**Supplemental Table 3.** Top 50 genes that are differentially expressed in pDCs of IFN$^{high}$ SLE patients in comparison with pDCs from healthy controls (HC).

| Gene | IFN$^{low}$ SLE pDCs vs. HC pDCs | | | IFN$^{high}$ SLE pDCs vs. HC pDCs | | |
|---|---|---|---|---|---|---|
| | Fold Change (log2) | *P* value | FDR | Fold Change (log2) | *P* value | FDR |
| ETV3 | 3.037 | < 0.001 | 0.017 | 2.959 | < 0.001 | 0.002 |
| ATF3 | 3.914 | < 0.001 | 0.052 | 5.209 | < 0.001 | 0.000 |
| LIN9 | -3.716 | < 0.001 | 0.058 | -2.423 | < 0.001 | 0.031 |
| LGALSL | 6.161 | < 0.001 | 0.007 | 6.240 | < 0.001 | < 0.001 |
| ZNF2 | -4.108 | < 0.001 | 0.058 | -2.907 | < 0.001 | 0.020 |
| LIPT1 | -3.224 | < 0.001 | 0.050 | -2.499 | < 0.001 | 0.002 |
| SEC24D | -2.559 | < 0.001 | 0.051 | -1.605 | 0.001 | 0.038 |
| CXCL2 | 6.379 | < 0.001 | < 0.001 | 4.163 | < 0.001 | 0.019 |
| CBR4 | -3.260 | < 0.001 | 0.052 | -1.546 | < 0.001 | 0.011 |
| DUSP1 | 2.903 | < 0.001 | 0.027 | 2.386 | < 0.001 | 0.021 |
| LTC4S | 5.509 | < 0.001 | 0.033 | 5.089 | < 0.001 | 0.003 |
| SNORD95 | 4.390 | < 0.001 | 0.007 | 3.106 | < 0.001 | 0.016 |
| PHF3 | 1.125 | < 0.001 | 0.015 | 1.531 | < 0.001 | 0.028 |
| LOC441242 | -2.798 | 0.001 | 0.068 | -2.494 | < 0.001 | 0.015 |
| FUT10 | -3.275 | 0.002 | 0.100 | -2.388 | < 0.001 | 0.016 |
| LOC102724580 | 3.699 | 0.001 | 0.088 | 4.080 | < 0.001 | 0.004 |
| CCL19 | 4.468 | < 0.001 | 0.057 | 3.828 | < 0.001 | 0.020 |
| CTSL | 6.460 | < 0.001 | 0.040 | 4.114 | < 0.001 | 0.020 |
| PUDP | -2.632 | < 0.001 | 0.058 | -2.752 | < 0.001 | 0.008 |
| ATP7A | -4.215 | < 0.001 | 0.035 | -2.853 | < 0.001 | 0.020 |
| SAT1 | 1.642 | < 0.001 | 0.036 | 1.805 | < 0.001 | 0.009 |
| MIR6087 | 2.376 | < 0.001 | 0.050 | 2.411 | < 0.001 | 0.028 |
| ZFP91 | 2.619 | < 0.001 | 0.046 | 2.147 | 0.001 | 0.043 |
| PAAF1 | -3.453 | 0.001 | 0.066 | -1.973 | < 0.001 | 0.033 |
| DUSP8 | 6.050 | 0.001 | 0.085 | 6.392 | < 0.001 | 0.006 |
| IRAK3 | 6.179 | < 0.001 | 0.045 | 4.152 | < 0.001 | 0.007 |
| PLEKHA8P1 | -4.947 | < 0.001 | 0.025 | -2.810 | < 0.001 | 0.009 |
| ACVR1B | 5.196 | 0.001 | 0.059 | 4.169 | < 0.001 | 0.017 |
| RN7SL1 | 2.867 | < 0.001 | 0.049 | 2.648 | < 0.001 | 0.034 |
| RN7SL2 | 3.107 | < 0.001 | 0.033 | 2.806 | < 0.001 | 0.021 |
| ATG14 | 2.520 | < 0.001 | 0.058 | 1.640 | < 0.001 | 0.028 |
| REREP3 | 3.757 | < 0.001 | 0.046 | 3.672 | < 0.001 | 0.019 |
| IRAIN | 5.493 | < 0.001 | 0.015 | 4.774 | < 0.001 | 0.004 |
| CDH1 | 3.795 | 0.001 | 0.061 | 2.895 | < 0.001 | 0.028 |
| SCARNA21 | 2.549 | 0.001 | 0.074 | 2.980 | 0.001 | 0.041 |
| C5AR1 | 6.724 | 0.002 | 0.098 | 5.210 | < 0.001 | 0.031 |
| SLC8A2 | 5.243 | < 0.001 | 0.023 | 5.644 | < 0.001 | < 0.001 |
| IER2 | 2.416 | < 0.001 | 0.050 | 2.552 | < 0.001 | 0.001 |
| DNAJB1 | 2.108 | < 0.001 | 0.015 | 4.097 | < 0.001 | 0.005 |
| NPAS1 | 5.880 | < 0.001 | 0.020 | 5.469 | < 0.001 | 0.002 |
| LOC100008589 | 2.695 | < 0.001 | 0.028 | 2.969 | < 0.001 | 0.006 |
| MIR3687-1 | 3.321 | 0.002 | 0.098 | 3.448 | < 0.001 | 0.007 |
| MIR3687-2 | 3.321 | 0.002 | 0.098 | 3.448 | < 0.001 | 0.007 |
| LOC100861532 | 2.839 | < 0.001 | 0.007 | 2.930 | < 0.001 | 0.005 |
| WRB | -2.306 | 0.001 | 0.068 | -2.027 | < 0.001 | 0.008 |
| **pDC-specific transcription factors** | | | | | | |
| E2-2 (TCF4) | 0.691 | 0.118 | 0.537 | 0.313 | 0.578 | 0.852 |
| SPIB | 0.657 | 0.279 | 0.719 | 0.235 | 0.744 | 0.922 |

**Supplemental Table 4.** Genes that are differentially expressed in pDCs of both IFN$^{low}$ and IFN$^{high}$ SLE patients in comparison with pDCs from healthy controls (HC).
